# Sbh1/Sec61β is a specific signal peptide receptor within the Sec61 channel

**DOI:** 10.1101/2024.03.12.584615

**Authors:** Lalitha Yadhanapudi, Mark Lommel, Pratiti Bhadra, Guido Barbieri, Richa Singh, Martin Jung, Volkhard Helms, Terese Eisgruber, Florian Stengel, Arne Boergel, Kim Remans, Karin Römisch

## Abstract

In eukaryotes, protein secretion begins with protein translocation through the universally conserved Sec61 channel into the endoplasmic reticulum (ER). Its β-subunit, Sbh1 in yeast, enhances ER import of proteins with specific suboptimal signal peptides by an unknown mechanism. The Sbh1 cytosolic N-terminus consists of an intrinsically disordered, non-conserved region (IDR) that has never been visualized in active channel structures, but is close to the translocating polypeptide in the cytosolic channel vestibule. The Sbh1/Sec61β N-terminal IDR is followed by structured 15 amino acids and its C-terminal transmembrane helix, both of which are conserved. We show here that the proline and adjacent conserved residues at the Sbh1 cytosolic/transmembrane domain interface form a hinge that positions the Sbh1 cytosolic domain across the channel vestibule and orients it with respect to the lateral gate. This orientation is critical for Sbh1-dependent protein insertion into the channel. Sbh1-dependence of Sec61 channel insertion is a function of the signal peptide of the respective secretory protein. By chemical crosslinking of purified cytosolic domains of Sbh1 and its paralog Sbh2 to synthetic signal peptides derived from their respective client proteins, we show that the cytosolic domains of Sbh1 and Sbh2 contain specific signal peptide binding sites. The position of the crosslinked residues suggests that signal peptide binding is mediated by the IDRs. We conclude that Sec61β homologs directly recognize signal peptides of their substrates and guide them into the Sec61 channel; they thus control entry of specific proteins into the secretory pathway.

## INTRODUCTION

About 30% of all proteins pass through the secretory pathway. In eukaryotes, protein secretion begins with signal peptide-mediated targeting to the endoplasmic reticulum (ER) [O’Keefe and High, 2020]. This leads to protein translocation through the universally conserved Sec61 channel into the ER lumen [O’Keefe and High, 2020]. Signal peptides are typically 20-30 amino acids long, and consist of a hydrophobic core flanked by an N-terminal region with a net positive charge and a polar C-terminal region containing the signal peptide cleavage site [O’Keefe and High, 2020; von Heijne, 1985]. The first transmembrane domain (TMD) of membrane proteins can also serve as an ER-targeting signal and is inserted into the membrane by lateral opening of the Sec61 channel [O’Keefe and High, 2020]. During co-translational translocation, signal recognition particle (SRP) binds to the signal peptide or first TMD as they exit the ribosome and targets the ribosome-nascent chain complex to the ER membrane. There SRP binds to its receptor and the signal peptide or TMD subsequently inserts into the lateral gate of the Sec61 channel [Cross et al., 2009]. Post-translational translocation of secretory protein precursors with less hydrophobic signal peptides requires auxiliary components. In yeast, this takes place through the heptameric Sec complex composed of the Sec63 complex (Sec62, Sec63, Sec71, Sec72) and the Sec61 channel [Allen et al., 2019; Itskanov and Park, 2019; Wu et al., 2019].

The Sec61 channel is heterotrimeric, consisting of the pore-forming Sec61 subunit, and two peripheral subunits, Sbh1, and Sss1 in yeast (Sec61α, Sec61β, and Sec61γ in mammals) [Mandon et al., 2013]. The *SEC61* and *SSS1* genes are essential in yeast [Mandon et al., 2013]. Sec61/Sec61α comprises 10 transmembrane (TM) helices arranged around a central hydrophilic pore that is constricted by a ring of hydrophobic amino acids and occluded by a plug helix on the lumenal side [Voorhees and Hegde, 2016]. The Sec61 N-terminal amphipathic helix is deeply embedded in the membrane and essential for protein import into the ER [Elia et al., 2019]. Soluble proteins are translocated into the ER through the central pore by displacing the plug [Voorhees and Hegde, 2016]. Membrane proteins are integrated into the ER membrane by lateral opening of the Sec61 channel between TM helices 2/3 and 7/8 (lateral gate) [Voorhees and Hegde, 2016]. Sss1/ Sec61γ stabilizes Sec61 by forming a clamp around its C-terminal half [Esnault et al., 1994; Voorhees and Hegde, 2016].

*S. cerevisiae* contains a paralog of the Sec61 channel, the Ssh1 channel, which consists of the channel-forming Ssh1 protein, Sss1, and the Sbh1 paralog Sbh2 [Finke et al., 1996; Toikkanen et al., 1996]. The Ssh1 channel is non-essential, translocates exclusively co-translationally, and has a different client spectrum from the Sec61 channel [Finke et al. 1996; Wilkinson et al. 2001; Spiller and Stirling, 2011; Cohen et al., 2023].

Sbh1/Sec61β is peripherally associated with Sec61 and is tail-anchored [Mandon et al., 2013]. *S. cerevisiae* is temperature-sensitive when the *SBH1* gene and its paralog *SBH2* are simultaneously deleted [Finke et al., 1996; Toikkanen et al., 1996; Feng et al., 2007]. This defect is rescued when the cells are complemented with a truncated Sbh1 or Sbh2 consisting of the TMD and the first 5 amino acids of the cytosolic domain [Feng et al., 2007; Leroux and Rokeach, 2008]. The conserved TMD of Sbh1 interacts with TM1 and TM4 of Sec61 and with TM3 of Sec63 in the Sec complex [Zhao and Jäntti, 2009; Wu et al., 2019)]. Despite its central position suited to connect the Sec61 channel with the Sec63 complex, Sbh1 does not contribute to the stability of the Sec complex [Bhadra et al., 2021]. Its absence also only marginally affects general posttranslational protein import into the ER [Finke et al., 1996; Feng et al., 2007].

Mammalian Sec61β binds to ribosomes, and both Sec61β and Sbh1 mediate the interaction between signal peptidase and the Sec61 channel during co-translational translocation, resulting in efficient insertion of the ribosome-bound nascent chain into the Sec61 channel [Levy et al., 2001; Kalies et al., 1998; Antonin et al., 2000]. The Sbh1 paralog Sbh2 is critical for the association of SRP receptor with the Ssh1 channel [Jiang et al., 2008]. The IDR of the Sec61β cytosolic domain can be crosslinked to stalled ER-targeting sequences in the cytosolic vestibule of the mammalian Sec61 channel, but absence of Sec61β only has a modest effect on general co-translational import through the mammalian Sec61 channel [Laird and High, 1997; Kalies et al., 1998; Feng et al., 2007]. In the pathogenic yeast *Cryptococcus neoformans*, Sbh1 is not required for vegetative growth, but specifically controls the biogenesis of secretory virulence factors [Santiago-Tirado et al., 2023]. In *S. cerevisiae*, about 12% of signal peptides rely on Sbh1/Sec61β for efficient translocation into the ER [Barbieri et al., 2023]. Signal peptides of Sbh1-dependent substrates have lower hydrophobicity and often an inverse, or no charge bias and thus are suboptimal for insertion into the lateral gate of Sec61 [Barbieri et al., 2023]. Secretory virulence factors in *C. neoforman*s also have suboptimal signal peptides [Santiago-Tirado et al., 2023]. For a specific subset of Sbh1-dependent translocation substrates, Sbh1 N-terminal phosphorylation at S3 and T5 controls the amount of entry of these proteins into the ER [Barbieri et al., 2023]. These substrates include the ER-resident enzymes mannosidase 1 and glucosidase 1 whose concentration in the ER is tightly regulated and critical for ER proteostasis, and the yeast Translocon-Associated Protein complex alpha subunit, Irc22, whose function in yeast remains unexplored [Feng et al., 2007; Hosokawa et al., 2003; Barbieri et al., 2023; Lewis et al., 2024].

The role of the Sbh1/Sec61β cytosolic region is not understood. Its IDR has never been visualized in active Sec61 channels, so despite its proximity to the translocating chain, how it contributes to ER protein import remains unknown. While we have shown that phosphorylation of proline-flanked S3/T5 at the N-terminus of the Sbh1 IDR controls the amount of ER import for specific proteins, how this works mechanistically is unclear [Barbieri et al., 2023]. The function of the approximately 15 amino acids long, universally conserved membrane proximal domain of Sbh1/Sec61β is also unclear [Kinch et al., 2002].

We show here that the conserved hinge at the Sbh1 cytosolic/transmembrane domain interface is critical for orientation of the Sbh1 cytosolic domain across the Sec61 channel vestibule and for controlling ER import. We demonstrate that the Sbh1 and Sbh2 IDRs constitute specific signal peptide binding sites. We find that Sbh1 S3/T5 phosphorylation does not control signal peptide binding, but in yeast ER membranes results in an SDS-resistant conformational change of the Sbh1 cytosolic domain encompassing at least the first 23 amino acids. Taken together our data demonstrate that Sbh1 and Sbh2 are specific signal peptide receptors which orchestrate the insertion of specific subsets of secretory proteins into the Sec61 channel.

## RESULTS

### MD-Modelling of contacts between the cytosolic N-termini of Sbh1 and Sec61

We have shown previously that the N-terminal amphipathic helix of Sec61 is critical for protein import into the ER [Elia et al., 2019]. This helix is deeply embedded in the ER bilayer and close to the conserved region of the Sbh1 cytosolic domain [Wu et al., 2019]. We therefore asked whether these regions are physically contacting each other in the Sec61 channel. As detailed in the Methods section, we performed three independent MD simulations to identify putative contacts between the cytosolic N-termini of Sec61 and Sbh1.

The N-terminal portion of Sec61 was anchored to the lipid bilayer via its TMD1. Both N-termini were modelled into alpha-helical conformations and MD simulations were all started from contacting conformations, but with different angles between the helical axes of the N-termini. In the simulation started from an inter-helical angle of +30°, the segment ^19^EBIAPERK^26^ of Sec61 remained consistently in close contact (ca. 5 Å distance) to segment ^45^DEATG^49^ of Sbh1. The final snapshot of this simulation after 50 ns is shown in Fig. 1A, with the putative contact interface highlighted. The other two MD simulations did not show atomic contacts between the N-termini of Sec61 and Sbh1 within a 5 Å distance.

**Figure 1.**
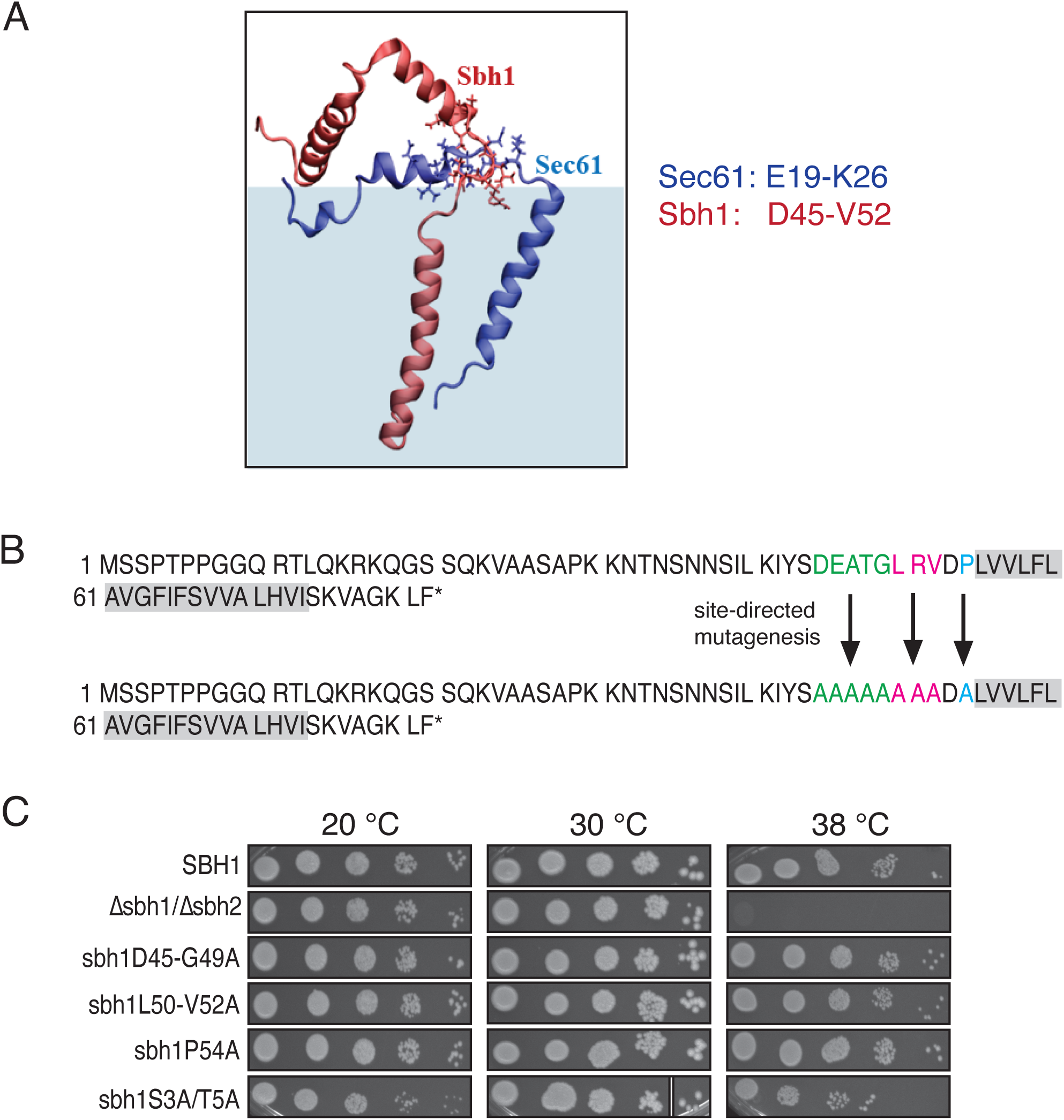
Identification of contacts between Sec61 N-terminal helix and the conserved section of the Sbh1 cytosolic domain by MD modelling. *A*, Final conformation of the N-termini of Sec61 (blue) and Sbh1 (red) after a 50 ns Molecular Dynamics simulation as described in Results. *B*, Site-directed mutagenesis of *SBH1* was used to generate the following mutants: *sbh1D45-G49A* (green), *sbh1L50-V52A* (red), and *sbh1P54A* (blue). *C*, Serial dilutions from 10^5^-10^1^ cells of wildtype and indicated *sbh1* mutants were grown on solid media at the indicated temperatures for 3 days. The spot corresponding to 10^1^ cells *sbh1S3A/T5A* on the 30 °C plate was spliced from the same plate to maintain its alignment with the remainder of the row. The *Δsbh1Δsbh2* double mutant and *sbh1S3A/T5A* served as controls.

### Sbh1 conserved domain mutants specifically affect ER-import of an Sbh1-dependent substrate

The putative contact site of Sbh1 with the Sec61 N-terminus was located in its conserved membrane-proximal region (aa 39-54, Fig. 1A). We generated three *sbh1* mutants in this region, by substituting either D45-G49 (green in Fig. 1B) or L50-V52 (pink in Fig. 1B) with A by site-directed mutagenesis. We also mutated the conserved residue P54 at the interface of the Sbh1 cytosolic domain and its membrane domain to A (blue in Fig. 1B). This residue interacts with TM4 of Sec61 and due to the rigidity of the prolyl-peptide bond orients the entire Sbh1 cytosolic domain with respect to the Sec61 channel [Zhao & Jäntti, 2009]. In the absence of functional *SBH1*, yeast are temperature-sensitive for growth at 38°C [Feng et al., 2007]. Many *sec61* mutants are also cold-sensitive [Pilon et al., 1998]. The reported growth defects are due to ER protein import defects of the mutant Sec61 channels which result in a weakened cell wall [Feng et al., 2007; Pilon et al., 1998; Barbieri et al., 2023]. We assayed the growth of our *sbh1* conserved domain mutants at permissive (30 °C) and restrictive (20° and 38 °C) temperatures. We did not observe temperature- or cold-sensitivity of growth of any of our *sbh1* mutants compared to wildtype, suggesting that they had no general ER protein import defects (Fig. 1C).

We then directly investigated ER import of specific proteins in our *sbh1* conserved domain mutants. We monitored the amounts of Sbh1 protein produced by each of the mutant strains and whether the mutations affected import of the Sbh1-dependent substrate glucosidase 1 (Gls1) into the ER [Feng et al., 2007; Barbieri et al., 2023]. We found that although all mutants and the wildtype were expressed from an identical plasmid and promoter, the *sbh1D45-G49A* mutant produced 1.5 times more Sbh1 than the wildtype (Fig. 2A, left panel). This resulted in Gls1 levels in the ER close to wildtype (Fig. 2A, right panel). In contrast, the *sbh1L50-V52A* mutant produced only 25% of wildtype Sbh1 protein, but had a 1.4x increased amount of Gls1 in the ER (Fig. 2A). The *sbh1P54A* mutant produced 75% of wildtype amounts of mutant Sbh1, but entry of Gls1 into the ER was similarly increased (1.6x, Fig. 2A). For these experiments it was critical to use freshly transformed cells as all mutants adapted over time such that the Gls1 levels in the ER became close to wildtype. Adaptation of growth and concomitant loss of translocation defects is common in yeast and have also been observed in other mutants affecting ER protein import [Mutka et al., 2001; Wilkinson et al., 2001; Jiang et al., 2008]. Our results suggest that within the Sbh1 conserved cytosolic region mutations in L50-V52 and P54 affect the gate-keeper function of Sbh1, partially alleviating the Sbh1-restricted access of Gls1 to the ER. Mutation of the D45-G49 region to A on the other hand seems to reduce the efficiency of Gls1 ER-import which the cell compensates by increasing Sbh1 mutant protein production.

**Figure 2.**
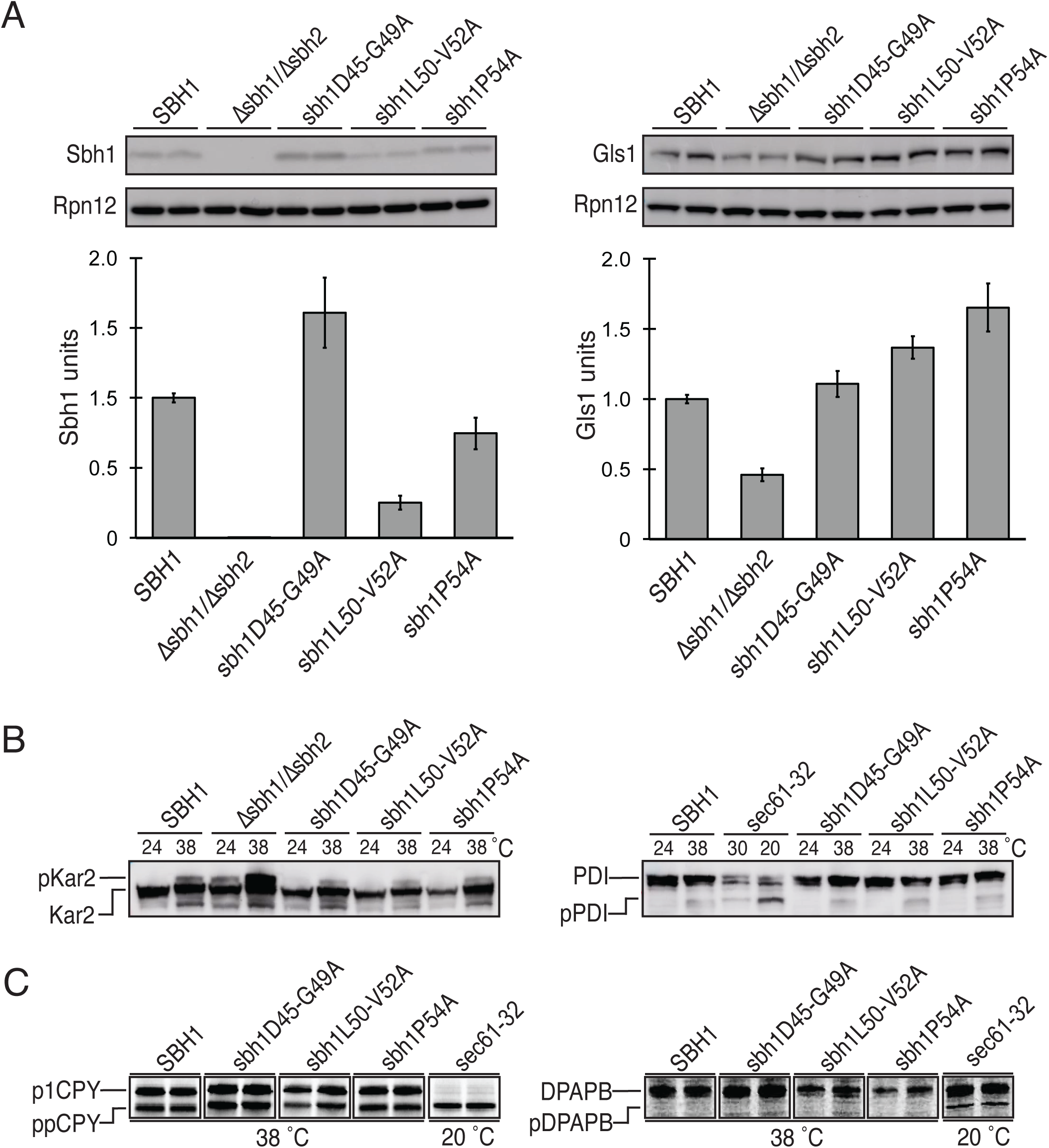
Mutations in the conserved cytosolic section of Sbh1 affect ER-import of specific substrates. *A*, Western blot analysis to investigate Sbh1 expression and glucosidase1 (Gls1) amounts in the ER in *sbh1* mutants. Rpn12 was used as the loading control. Wildtype *SBH1* and all mutants were expressed from a CEN plasmid in a *Δsbh1 Δsbh2* strain. Freshly transformed cells were grown to early exponential phase, lysed, proteins resolved by SDS-PAGE, transfered to nitrocellulose and proteins detected with specific antisera. Representative blots are shown on top. Graphs: Sbh1 and Gls1 amounts in 7 randomly picked colonies of each *sbh1* mutant from 3 independent experiments. Signals from Western Blots with the indicated antibodies were quantified and shown relative to their levels in *SBH1* wildtype cells. *B*, Western blot analysis showing the translocation of cytosolic precursors of Kar2 and PDI at permissive (24°C) and restrictive (38°C) temperatures. The permissive temperature of 24°C was chosen to reduce adaptation of the mutants. Cells were grown to early exponential phase and lysed, proteins resolved by SDS-PAGE and transferred to nitrocellulose, and proteins detected with specific antisera. The *Δsbh1Δsbh2* mutant was used control for pKar2 accumulation. For pPDI accumulation *sec61-32* was used as control, grown at 30°C (permissive) and shifted to 20°C (restrictive) for 3 h. *C*, Secretory precursor accumulation test by pulse-labelling of cells in early exponential phase with [^35^S]-Met/Cys for 5 min (ppCPY) or 15 min (pDPAPB). After the pulse, cells were lysed and proteins immunoprecipitated with specific antisera. Proteins were resolved by SDS-PAGE and detected by autoradiography. Positive control used for both experiments was *sec61-32*.

To investigate whether our *sbh1* mutants also affected general protein import into the ER, we investigated ER import of the precursors of Kar2 (BiP), protein disulfide isomerase (Pdi1), carboxypeptidase Y (CPY), and dipeptidyl aminopeptidase B (DPAPB) at the restrictive temperature 38 °C. Kar2 is dependent on Sbh1 for its import, but independent of Sbh1 S3/T5 phosphorylation [Barbieri et al., 2023]. Kar2 import remained unaffected in our *sbh1* mutants (Fig. 2B, left). We also did not detect any significant cytosolic precursor accumulation of post-translationally translocated Pdi1 in our *sbh1* mutants as compared to the wildtype (Fig. 2B). The *sbh1* cytosolic conserved domain mutants also efficiently translocated post-translationally imported prepro-CPY (Fig. 2C, left). Co-translational import of pDPAPB was also not affected (Fig. 2C, right). Taken together, our results suggest that mutations in the conserved section of the Sbh1 cytosolic region do not generally affect co- or posttranslational ER import, but that their translocation phenotypes are restricted to proteins that depend on N-terminally S3/T5-phosphorylated Sbh1 for their import into the ER.

In order to better understand the effects of the mutations in the conserved part of the Sbh1 cytosolic region on import of an Sbh1-dependent translocation substrate, we subjected Sbh1 wildtype and *sbh1P54A*, *sbh1L50-V52A*, and *sbh1D45-G49A* mutants to Rosetta *ab initio* modelling [Alford et al., 2017]. The resulting structures are shown in Supplementary Figure 1 with the Sbh1 C-terminal transmembrane helix in the same fixed position - determined by the side chains of P54 and V57 which are known to contact TM4 of Sec61 -, such that changes in the orientation of the cytosolic domain due to the mutations are evident [Zhao and Jäntti, 2009]. In wildtype Sbh1, P54 orients the Sbh1 cytosolic domain across the vestibule of the Sec61 channel, with the N-terminus facing up towards the ribosome (Supp. Fig. 1, top left). This orientation is similar to that of Sbh1 in a recent cryo-EM structure of the closed Sec61 channel of the thermophilic yeast *Chaetomium thermophilum* [Yang et al., 2025]. In contrast, in the *sbh1P54A* mutant the Sbh1 cytosolic domain is oriented towards the opposite side, away from the channel (Supp. Fig. 1, top right). Similarly, in *sbh1L50-V52A*, the cytosolic domain is no longer positioned across the Sec61 channel vestibule, but oriented up and away from the channel (Supp. Fig. 1, bottom right). The structure predictions for these mutants suggest that the Sbh1 cytosolic domain has a barrier function and physically blocks access to the channel for Gls1 in the Sbh1 wildtype, but not in the hinge region mutants *sbh1P54A* and *sbh1L50-V52A*. Rosetta ab initio modelling of the *sbh1D45-G49A* showed a more subtle change: here, the cytosolic domain was still positioned across the channel vestibule, but its orientation was changed, with the N-terminus now pointing towards the channel (Supp. Fig. 1, bottom left). Our data on Gls1 import suggest that in this orientation, the barrier function is still intact, but the changed orientation likely makes Sbh1 less efficient in promoting ER import of Gls1 which the cells expressing the *sbh1D45-G49A* mutant compensate by upregulating mutant protein production (Fig. 2A). In conclusion, these structural models suggest that the orientation of the Sbh1 cytosolic domain with respect to the Sec61 channel is critical for controlling ER import of substrate proteins dependent on S3/T5-phosphorylated Sbh1.

### The Sbh1 cytosolic domain binds directly to specific signal peptides

We have shown above that the Sbh1 cytosolic domain orientation is critical for function and we have shown previously that Sbh1-dependence of Sec61 channel insertion of secretory proteins is a function of their signal peptides [Barbieri et al., 2023]. We therefore asked whether the Sbh1 cytosolic domain can discriminate between signal peptides of its substrates and those of non-substrate proteins and whether it can directly bind to signal peptides. We purified the overexpressed Sbh1 cytosolic domain from *E. coli* as in Materials & Methods and incubated it for 30 min at 20°C with 10x excess synthetic biotinylated signal peptides derived from proteins dependent on Sbh1 (Irc22, Gls1, Mns1) and from Sbh1-independent secretory proteins (carboxypeptidase Y, CPY, and prepro alpha factor, ppaF) [Barbieri et al., 2023]. Complexes were crosslinked with the amino-reactive crosslinker disuccinimidyl suberate (DSS). After quenching, proteins were resolved on Tricine gels, transferred to nitrocellulose and detected with either antibodies against Sbh1 or against biotin. The Sbh1 cytosolic domain could only be crosslinked to signal peptides derived from Sbh1-dependent translocation substrates, suggesting that Sbh1 can discriminate between signal peptides of Sbh1-dependent substrates and signal peptides from non-substrates, and that it directly binds to these signal peptides (Fig. 3A).

**Figure 3.**
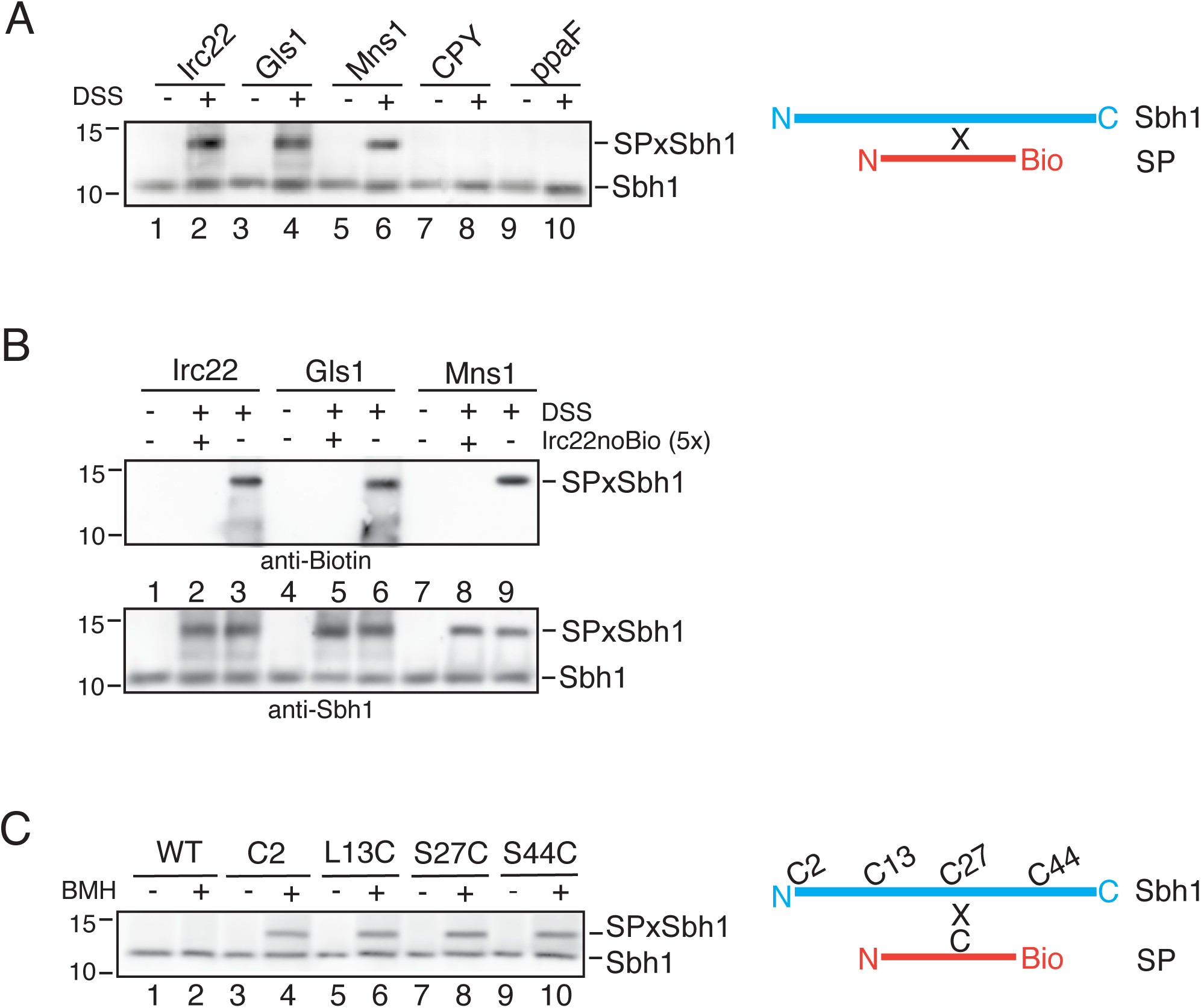
The Sbh1 cytosolic domain specifically recognizes signal peptides. In all experiments, overexpressed purified cytosolic domains of Sec61β homologs (15 µM) and synthetic, biotinylated signal peptides (150µM) were incubated in the absence or presence of 150 µM crosslinker at 20 °C for 30 min. Reactions were quenched, samples resolved on Tricine gels, and proteins transferred to nitrocellulose. The Sbh1 cytosolic domain and its crosslinking products were detected by immunoblotting with the Sbh1 (39-48) antibody. Signal peptide crosslinking products were also detected with an anti-Biotin antibody. Molecular weight in kDa is indicated on the left of each panel. All experiments were repeated at least twice. *A*, The Sbh1 cytosolic domain was incubated with signal peptides from Sbh1-dependent (Irc22, Gls1, Mns1) and Sbh1-independent (CPY, ppaF) translocation substrates and the absence or presence of the amino-reactive crosslinker disuccinimidyl suberate (DSS). After gel electrophoresis and transfer to nitrocellulose, proteins were detected with anti-Sbh1 antibodies. *B*, The Sbh1 cytosolic domain was incubated with biotinylated Sbh1-dependent signal peptides (Irc22, Gls1, Mns1) and DSS in the absence or presence of a 5-fold excess of non-biotinylated Irc22 signal peptide for competition as indicated. After gel electrophoresis and transfer to nitrocellulose, signal peptides crosslinked to Sbh1 were detected with anti-Biotin (upper panel) and all Sbh1 and crosslinking products were detected with anti-Sbh1 antibodies (lower panel). *C*, Purified wildtype Sbh1 cytosolic domain and point mutants with unique cysteine residues introduced at the indicated positions were incubated with the Irc22 signal peptide which naturally contains a unique cysteine. Samples were crosslinked with the sulfhydryl-reactive crosslinker bismaleimidohexane (BMH). After gel electrophoresis and transfer to nitrocellulose, proteins were detected with anti-Sbh1 antibodies.

We next asked whether all crosslinkable signal peptides (Irc22, Gls1, Mns1) occupied the same binding site in the Sbh1 cytosolic domain. We incubated the Sbh1 cytosolic domain with the three substrate signal peptides as above, but included a second sample for each to which we added a 5-fold excess of unbiotinylated Irc22 signal peptide. After crosslinking and quenching, samples were resolved on Tricine gels, transferred to nitrocellulose and crosslinked signal peptides were detected by immunoblotting with anti-biotin antibodies (Fig. 3B, upper) and Sbh1 and signal peptide/Sbh1 complexes with anti-Sbh1 antibodies (Fig. 3B, lower). As expected, excess unbiotinylated Irc22 signal peptide competed with crosslinking biotinylated Irc22 signal peptide to Sbh1 (Fig. 3B, compare lanes 3 with 2, upper panel). In addition, excess Irc22 signal peptide was also able to compete with binding of biotinylated Gls1 and Mns1 signal peptides to Sbh1 (Fig. 3B, compare lanes 6 with 5, and 9 with 8, upper panel). In each case, unbiotinylated Irc22 signal peptide also resulted in a crosslinked band detectable with the Sbh1 antibody (Fig. 3B, lanes 2, 5, 8, lower panel). We conclude that the Sbh1 cytosolic domain forms a single binding site to bind all Sbh1-dependent signal peptides.

To measure the extent of contact between signal peptides and the Sbh1 cytosolic domain we introduced unique cysteine residues at different positions into Sbh1 as shown in Fig. 3C, purified the overexpressed cysteine variants from *E. coli,* and used them in crosslinking experiments with the Irc22 signal peptide, which naturally contains a unique cysteine, with the sulfhydryl-reactive homobifunctional crosslinker bismaleimidohexane (BMH). C2 was inserted after the starting methionine of Sbh1. In all other variants, individual amino acids were replaced by cysteines at the positions as indicated in Fig. 3C. Human Sec61β naturally contains a unique cysteine residue at a position approximately equivalent to S27 in Sbh1 which has been found crosslinked to stalled ER import substrates [Laird and High, 1997; MacKinnon et al., 2014; Jadhav et al., 2015]. As shown in Fig. 3C, the cysteine in the Irc22 signal peptide could be crosslinked to each Sbh1 cysteine variant, but not to wildtype Sbh1 which does not contain cysteine (Fig. 3C, compare lane 2 to lanes 4, 6, 8, 10). This suggests that interaction of a signal peptide with Sbh1 can be initiated from any position along its cytosolic domain.

To identify the natural positions of signal peptide crosslinks in the Sbh1 cytosolic domain, we generated scaled-up DSS crosslinking reactions of Sbh1 with the Irc22, Mns1, or Gls1 signal peptide and analyzed the resulting complexes by crosslinking-mass spectrometry [Leitner et al., 2014]. We identified crosslinks between K2 of the Mns1 signal peptide and K23 of Sbh1 (Supp. Fig. 2), as well as crosslinks between K5 of the Gls1 signal peptide and K17, K30, and K31 within the IDR of Sbh1, spanning residues 1-38 (Supp. Fig. 2). The Gls1 signal peptide contains additional K residues at positions 7 and 10 which were not found to be crosslinked (Supp. Fig. 2). K41 within the conserved region of the Sbh1 cytosolic domain was also not observed to crosslink with any of the signal peptides (Supp. Fig. 2). As the Irc22 signal peptide lacks lysine residues, DSS-mediated crosslinking to Sbh1 could only have occurred via its N-terminus; such N-terminal crosslinks, however, were not detected by our mass spectrometry analysis (Supp Fig. 2). Taken together, our data suggests that signal peptide interactions with Sbh1 take place primarily in the IDR of its cytosolic domain.

### In the yeast ER Sbh1 S3/T5 phosphorylation results in a conformational change of the N-terminal half of the Sbh1 cytosolic domain

We have shown that phosphorylation of Sbh1 at S3 and T5 is critical for the amount of Sbh1-dependent import of Gls1, Mns1, and Irc22 precursors into the ER [Barbieri et al., 2023]. The Sbh1 cytosolic domains we used for signal peptide crosslinking had been purified from *E. coli*, and were therefore not phosphorylated. Our crosslinking results shown in Fig. 3 demonstrate that S3/T5 phosphorylation is not required for Irc22, Gls1, or Mns1 signal peptide binding to Sbh1. To investigate whether there were any effects of S3 and T5 phosphorylation on signal peptide interaction with Sbh1, we chemically synthesized the Sbh1 cytosolic domain with either S3 or T5 or both residues phosphorylated, and used these synthetic cytosolic domains in signal peptide crosslinking experiments. As shown in Fig. 4A, phosphorylation of either or both residues had no or only a minor effect on interaction with the Gls1 signal peptide (top panel), whereas only the T5-phosphorylated synthetic Sbh1 cytosolic domain could interact with the Irc22 signal peptide (middle panel), and none of the phosphorylated forms of the Sbh1 cytosolic domain could be crosslinked to the Mns1 signal peptide (bottom panel). Since all three proteins, Gls1, Irc22, and Mns1, require both phosphorylatable S3 and T5 residues in Sbh1 for efficient ER import in yeast cells [Barbieri et al., 2023], our data suggest that phosphorylation at S3 and T5 does not primarily regulate signal peptide binding to Sbh1, but a different step or interaction in the ER translocation process.

**Figure 4.**
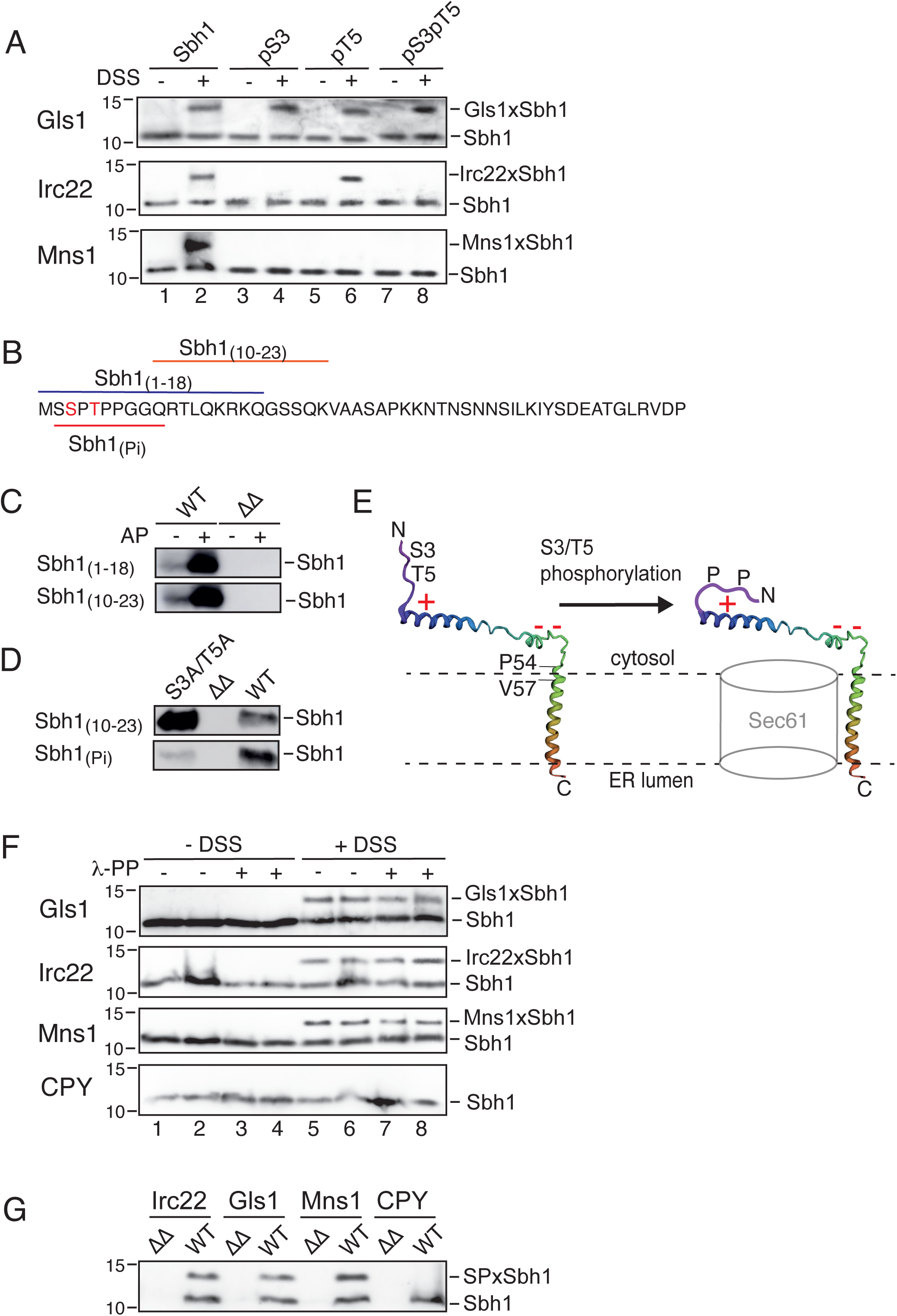
Sbh1 S3/T5 phosphorylation causes a conformational change in Sbh1. **A)** Unphosphorylated, S3, T5, or S3/T5 phosphorylated Sbh1 cytosolic domain was incubated with the indicated synthetic signal peptides and amino-specific DSS crosslinker as described in Fig. 3A. Crosslinked samples were resolved on Tricine gels, transferred to nitrocellulose, and Sbh1 and Sbh-SP complexes detected with antibodies to Sbh1. **B)** Schematic representation of peptide antibody epitopes in Sbh1. Sbh1(1-18) (blue) was raised against first 18 residues of Sbh1, Sbh1(10-23) (orange) raised against residues 10 to 23 of Sbh1, and Sbh1(Pi) (red) was raised against residues 2 to 10 and with S2-acetylation and S3-phosphorylation (’anti-phospho-Sbh1’). **C)** Antibody recognition test for Sbh1(1-18) and Sbh1(10-23) antibodies. Equal amounts of microsomal protein (50 µg) from *SBH1 SBH2* wildtype (WT) and *Δsbh1Δsbh2* (ΔΔ) strains were treated (+) with alkaline phosphatase (AP) to dephosphorylate microsomal proteins exposed to the cytosolic face or left untreated (-). Proteins were resolved by SDS-PAGE, transferred to nitrocellulose, and probed with the indicated antibodies. **D)** Antibody recognition test for Sbh1(10-23) and Sbh1(Pi) antibodies. Microsomal protein (50 µg) from *SBH1 SBH2* wildtype (WT), *Δsbh1Δsbh2* (ΔΔ), and *sbh1S3A/T5A* (S3A/T5A) were resolved on SDS-PAGE and transferred to nitrocellulose and probed with the indicated antibodies. **E)** Model showing conformational change of Sbh1 upon S3/T5 phosphorylation. Sbh1 was folded *ab initio* using Rosetta software (Alford et al., 2017). Sbh1 orientation with respect to the Sec61 channel was determined using the interaction of P54 and V57 (indicated on left) with Sec61. The two negatively charged residues (D45 and E46) in the conserved region anchoring the Sbh1 cytosolic domain to the Sec61 channel vestibule are indicated by (- -), the conserved positively charged patch (K15-K17) by (+). In the S3/T5 unphosphorylated state (left) the epitopes for Sbh1 (1-18) and Sbh1 (10-23) antibodies are accessible. Upon Sbh1 S3/T5 phoshorylation (right), negatively charged phosphate groups (P) may bind to the conserved positive patch (+), thus occluding the epitopes for Sbh1 (1-18) and Sbh1 (10-23) antibodies. F) Signal peptide crosslinking to full length Sbh1 in yeast microsomes. Wildtype *SBH1 SBH2* yeast microsomes were phosphatase-treated or mock-treated as indicated, incubated with the indicated synthetic signal peptides and crosslinked with DSS. Proteins were resolved on Tricine gels, transferred to nitrocellulose, and Sbh1 and Sbh1 crosslinks detected with Sbh1 (39-48) antibody. G) Control experiment for F): Wildtype (WT) or *Δsbh1Δsbh2* mutant (ΔΔ) yeast microsomes were phosphatase-treated, incubated with the indicated signal peptides, and crosslinked with DSS and processed as above.

Phosphorylation can have large scale effects on IDR conformation [Bah et al., 2015]. We therefore investigated the conformation of full-length phosphorylated Sbh1 in the context of the Sec61 channel in the yeast ER membrane. We asked whether accessibility to N-terminal Sbh1 antibodies was altered by phosphorylation of S3 and T5 of Sbh1 in the ER. We isolated wildtype yeast ER membranes and mock-treated or phosphatase-treated equal amounts of membranes to remove phosphate residues from Sbh1. Samples were resolved by SDS-PAGE and Sbh1 detected with antibodies raised against Sbh1 residues 1-18 or Sbh1 residues 10-23 (Fig. 4B, C). Both antibodies recognized dephosphorylated Sbh1 substantially better than the phosphorylated form (Fig 4C). This suggests a large, SDS-resistant conformational change in the Sbh1 N-terminus upon S3/T5 phoshorylation that occludes the epitopes for the Sbh1 (1-18) antibody (which includes the S3 and T5 phosphorylation sites, Fig. 4B) and the Sbh1 (10-23) antibody (which does not include these phosphorylation sites, Fig. 4B). Our data also suggest that at steady state the vast majority of Sbh1 in the cell is S3/T5 phosphorylated (Fig. 4C; compare signals before and after phosphatase treatment).

To investigate whether the conformational change can be attributed to the S3/T5 phosphorylation, we compared Sbh1 (10-23) antibody reactivity of the Sbh1S3A/T5A mutant protein with that of the wildtype protein. Both proteins are stable and expressed in equal amounts [Barbieri et al., 2023]. We found that anti-Sbh1 (10-23) reactivity towards the Sbh1S3A/T5A mutant protein was substantially increased compared to wildtype Sbh1 (Fig. 4D, upper panel). For comparison, reactivity with the antibody raised against the phosphorylated Sbh1 N-terminus is shown (Fig 4B, Fig. 4D, lower panel). This antibody preferentially recognizes the N-terminally phosphorylated wildtype Sbh1 and reacts poorly with the Sbh1S3A/T5A mutant protein (Fig. 4D). Since the epitope for anti-Sbh1 (10-23) does not include the S3/T5 phosphorylation sites and wildtype Sbh1 is >90% phosphorylated at steady state, this experiment confirms that the phosphorylation of Sbh1 at S3 and T5 residues induces a conformational change at its N-terminus involving residues at least up to K23 (Fig 4D, E).

We then investigated whether the S3/T5-phosphorylation-induced conformational change in Sbh1 observed in ER membranes affected signal peptide binding. To address this question, we used microsomes from an *SBH1 SBH2* wildtype strain that had been treated with phosphatase or mock-treated, and incubated these membranes with Gls1, Irc22, or Mns1 signal peptide and DSS. As shown in Fig. 4F, Sbh1 in yeast microsomes could be crosslinked to the Gls1, Irc22, and Mns1 signal peptides regardless of whether the membranes had been phosphatase-treated or not. Nevertheless signal peptide interaction with phosphorylated wildtype Sbh1 in microsomes was still specific, as the CPY signal peptide could not be crosslinked to Sbh1 in ER membranes (Fig. 4F, bottom panel). We also confirmed that no Sbh1-related band were detectable in microsomes derived from a *Δsbh1 Δsbh2* strain (Fig. 4G). As the conformational change triggered by Sbh1 S3/T5 phosphorylation in the yeast ER had no effect on signal peptide binding to the Sbh1 IDR in the context of the Sec61 channel in the ER membrane, this confirms our conclusion that S3/T5 phosphorylation does not regulate signal peptide binding to Sbh1. It is therefore likely that Sbh1 S3/T5 phosphorylation regulates a different step in translocation or interaction with another Sec61 channel accessory protein.

### The IDRs of Sbh1 and Sbh2 are specific signal peptide binding sites

To investigate whether the identity of the Sbh1 IDR was critical for determining signal peptide binding specificity, we made use of the Sbh1 paralog Sbh2. Due to an ancient genome duplication, *S. cerevisiae* express a Sec61-homologous (Ssh1) channel, with an Sbh1 paralog, Sbh2 [Finke et al., 1996; Toikkanen et al., 1996]. The Ssh1 channel cannot substitute for the Sec61 channel, is not essential, and has a different substrate spectrum [Finke et al., 1996; Wilkinson et al., 2001; Cohen et al., 2023]. Substrate specificity is conferred by the Sbh2 subunit of the channel [Spiller and Stirling, 2011]. Sbh1 and Sbh2 are almost identical in their TMDs and conserved cytosolic regions, but differ in their IDRs (Fig. 5A). Sbh2 also lacks the N-terminal proline-flanked phosphorylation sites that are critical for controlling efficient ER import of Mns1, Gls1, and Irc22 (Fig. 5A) [Barbieri et al., 2023]. Our Sbh1 N-terminal antibody poorly recognizes Sbh2 (Supp. Fig. 3, upper panel), whereas a specific antibody against the Sbh2 N-terminal peptide does not recognize Sbh1 at all (Supp. Fig. 3, lower panel), suggesting that the N-terminal IDRs of Sbh1 and Sbh2 have different conformations in the ER.

**Figure 5.**
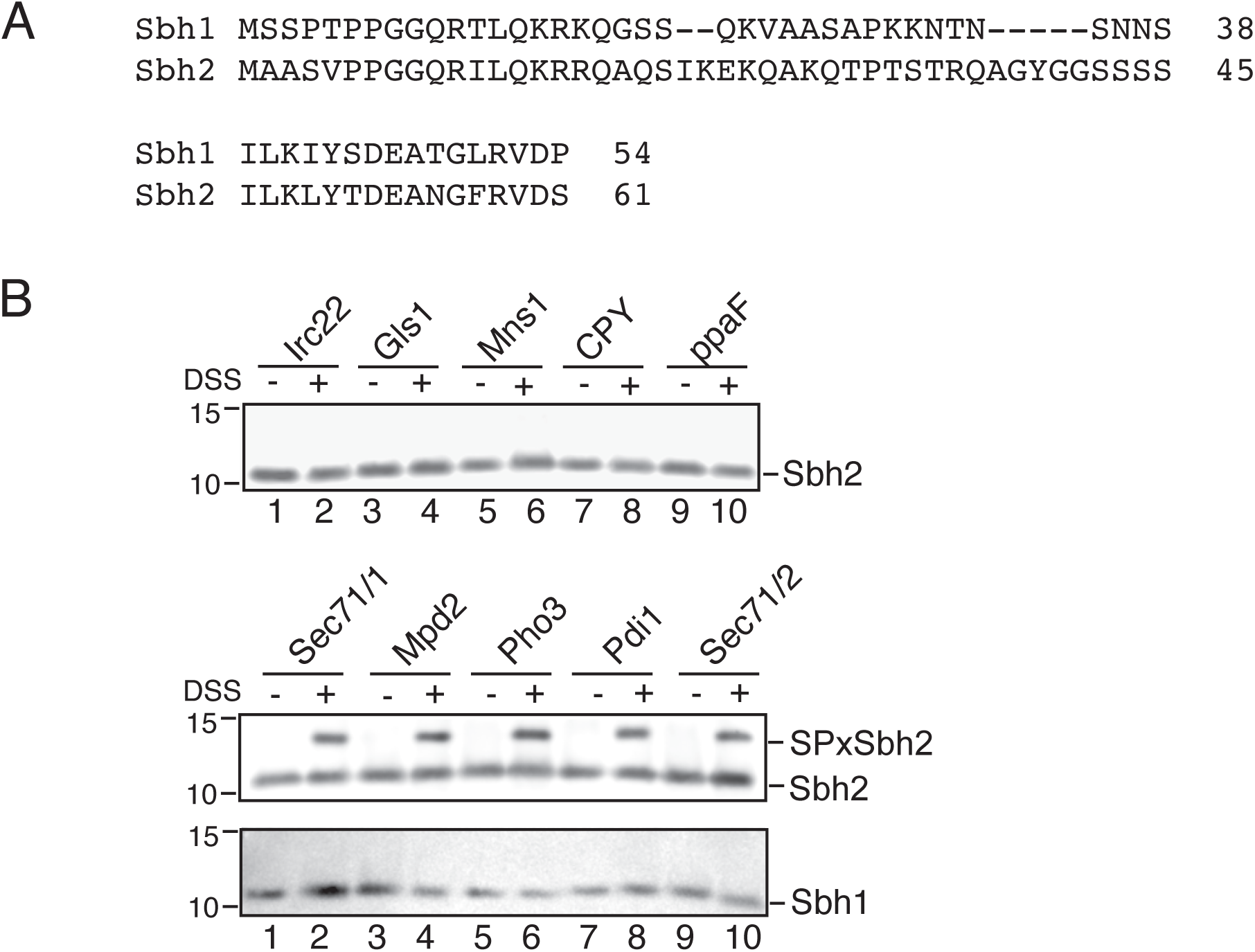
The Sbh2 cytosolic domain specifically recognizes and binds to signal peptides. *A*, Alignment of the Sbh1 and Sbh2 cytosolic domains. Top line shows the IDRs, bottom line the conserved parts of the cytosolic domains. *B*, The Sbh2 cytosolic domain was incubated with signal peptides derived from Sbh1-dependent proteins (Irc22, Gls1, Mns1) and Sbh1- independent proteins (CPY, ppaF) (top panel) or signal peptides from Sbh2-dependent proteins (Mpd2, Pho3, Pdi1) or a transmembrane ER targeting signal from an Sbh2-dependent protein (Sec71; two versions with different sections of adjacent residues) (middle panel). Crosslinking was done with DSS as indicated. As a control, Sbh2-dependent targeting peptides were also incubated with the Sbh1 cytosolic domain and crosslinked (bottom panel). After gel electrophoresis and transfer to nitrocellulose, proteins were detected with anti-Sbh1 (1-18) antibodies which cross-react with Sbh2 (Supp Fig. 3).

We overexpressed the Sbh2 cytosolic domain in *E. coli* and used the purified protein in signal peptide crosslinking experiments with DSS as in Fig. 3A. We were unable to detect any interactions of the Sbh2 cytosolic domain with signal peptides derived from Sbh1-dependent proteins (Irc22, Gls1, Mns1) or from Sbh1-independent proteins (CPY, ppaF) (Fig. 5B, top panel). In contrast, when we incubated the Sbh2 cytosolic domain with ER targeting peptides derived from Ssh1 channel-dependent substrate proteins (signal peptides: Mpd2, Pho3, Pdi1; TMD with different flanking regions: Sec71/1, Sec71/2), we found that all of these targeting peptides could interact with Sbh2 (Fig. 5B, middle panel) [Cohen et al., 2023; Spiller and Stirling, 2011; Wilkinson et al., 2001]. None of the Sbh2-interacting ER targeting peptides were able to bind to Sbh1 (Fig. 5B, bottom panel). In conclusion, our work demonstrates that within the Sec61 channel Sec61β homologs are highly specific receptors for select ER targeting signals.

## DISCUSSION

Sbh1/Sec61β is the only non-essential subunit of the universally conserved Sec61 channel [Finke et al., 1996; Toikkanen et al., 1996]. Although all organisms have this subunit, how it contributes to ER protein import remains unclear. In this paper, we address the function of its cytosolic domain which due to its intrinsic disorder has never been observed in the context of a translocating, active Sec61 channel.

We have shown here that the conserved section of the Sbh1 cytosolic domain - 15 amino acids proximal to the TMD - is required for positioning the Sbh1 cytosolic domain across the Sec61 channel vestibule and orienting its N-terminus towards the cytosol (Fig. 2, Supp. Fig. 1). Alanine mutations in the hinge region at the TMD which likely re-position the cytosolic domain away from the Sec61 channel (Supp. Fig. 2, right) led to increased ER import of the Sbh1-dependent substrate Gls1, suggesting that the Sbh1 cytosolic domain restricts and controls access to the Sec61 channel (Fig. 2A, right). This had also been suggested by Yang and colleagues, who recently solved the cryo-EM structure of a ribosome-associated Snd3 translocon complex, which includes the Sec61 channel in the closed conformation [Yang et al., 2025]. In this complex from the thermophilic yeast *Chaetomium thermophilum*, the N-terminus of Sbh1 is found in a similar orientation as in our Rosetta model (Supp Fig 1, top left) and bound to the ribosome [Yang et al., 2025].

Alanine mutations further N-terminal in the conserved part of the Sbh1 cytosolic domain (D45-G49) required overexpression of mutant Sbh1 to achieve wildtype amounts of Gls1 import (Fig. 2A). Rosetta modelling suggested a re-orientation of the cytosolic domain in this mutant with the N-terminus facing towards the ER membrane (Supp. Fig 2, lower left) [Das et al., 2008]. The *C. thermophilum* equivalents of two of the mutated residues (D45 and E46) have been shown to anchor the Sbh1 cytosolic domain to TMD3 of the Sec61 channel and to form an intramolecular salt bridge with the conserved, positively charged amino acid 5 positions further C-terminal (R51 in *S. cerevisiae* Sbh1). The conservation of these interactions and our experimental results suggest that the orientation of the Sbh1 cytosolic domain across the channel and with the N-terminus towards the cytosol is critical for function. The Sbh1 N-terminus in the *C. thermophilum* structure is bound to the ribosome [Yang et al., 2025]. The Sbh1-dependent ER important substrates we identified are likely imported co-translationally [Barbieri et al., 2023], so the *S. cerevisiae* Sbh1 N-terminus may also be in contact with the ribosome during their ER import.

When the Sec61 channel opens, the N-terminal half of Sec61 swings away from the C-terminal half by about 20 degrees [Voorhees and Hegde, 2016]. As the conserved sections of Sbh1 (TMD and the first 15 cytosolic amino acids, I38-I74 in *S. cerevisiae*) contact TMD1, 3, and 4 of Sec61, they have the capacity to stabilize the channel in the open conformation and hence increase the window of time for insertion of the Sbh1-dependent suboptimal signal peptides [Zhao and Jäntti, 2009; Yang et al., 2025; Barbieri et al., 2023]. This may also explain how the Sbh1 and Sbh2 TMDs with only 5 cytosolic amino acids can complement the temperature-sensitivity of a *Δsbh1 Δsbh2* yeast strain and that some organisms survive with a Sec61β/Sbh1 that lacks the IDR and only consists of the conserved part of the cytosolic section attached to a TMD [Feng et al., 2007; Leroux and Rokeach, 2008; Kinch et al., 2002].

In bacteria, the Sec61 homologous SecYEG channel exports proteins directly across the plasma membrane. The Sbh1 homolog SecG stabilizes the interaction of mutant SecY forms in the open conformation with SecE (the Sss1/Sec61g homolog) [Belin et al., 2015]. Like Sbh1, SecG restricts access of specific signal peptides to the SecY channel, and promotes export of proteins with suboptimal mutant signal peptides [Belin et al., 2015].

We show here that the IDRs of Sbh1 and Sbh2 bind signal peptides with different and high specificity (Fig. 3, Fig. 5, Supp Fig 2). Although contact with ER targeting sequences of mammalian Sec61β had been observed before, this was with stalled translocation substrates and hence thought to occur due to the increased residence time of these targeting sequences in the channel vestibule [Laird and High, 1997; MacKinnon et al., 2014; Jadhav et al., 2015]. A specific and functionally critical interaction between the IDRs of Sec61β homologs and ER targeting sequences has not been observed before. We have shown previously that this interaction allows the controlled import of a subset of ER translocation substrates whose targeting peptides are suboptimal for insertion into the Sec61 channel [Barbieri et al., 2023]. It is therefore likely that the targeting sequences of the proteins that have been used in translocation and crosslinking experiments with mammalian ER in the past, and which had been chosen for optimal translocation efficiency, were not dependent on Sec61β for ER import. There are, however, examples of mammalian secretory proteins whose biogenesis relies on Sec61β [Besse et al., 2017]. We would predict that ER targeting peptides of these proteins directly interact with the IDR of Sec61β, and that this interaction is critical for the ER import of these proteins.

Efficient ER import of Gls1, Mns1, and Irc22 relies on phoshorylation of S3 and T5 in the Sbh1 N-terminus [Barbieri et al., 2023]. Particularly striking is the convergent evolution of the proline-flanked Sbh1/Sec61β T5 phosphorylation site in yeast and mammals, suggesting that this site has an important function [Soromani et al., 2012]. Proline-flanked phosphorylation sites are also present in Sbh1 of pathogenic yeast species, where Sbh1 controls biogenesis of secretory virulence factors, but are largely absent from non-pathogens [Wei et al., 2020; Santiago-Tirado et al., 2023]. We expected S3/T5 phosphorylation to control signal peptide binding to the Sbh1 cytosolic domain. Instead, we found that phosphorylation at these positions had differential effects on interaction of these three signal peptides with a synthetic Sbh1 cytosolic domain (Fig. 4A), and no effect on their interaction with intact Sbh1 in the context of the Sec61 channel in ER membranes (Fig. 4F). In full length Sbh1 in the yeast ER, the S3/T5 phosphorylation sites controlled a large, SDS-resistant conformational change involving at least half of the N-terminal IDR (Fig. 4C, D, E). In yeast ER membranes, about 90% of Sbh1 was found in this conformation at steady state (Fig. 4C). We conclude that the signal binding ability of Sbh1is not regulated by S3/T5 phosphorylation. The phosphorylation-induced conformational change in Sbh1 may instead control interaction with an as yet unidentified Sec61 channel accessory factor that controls the amount of Gls1, Mns1, and Irc22 import into the yeast ER. Sbh1 S3/T5 phosphorylation may also affect its interaction with the ribosome [Yang et al., 2025] and explain why the cytosolic domain of the SRP receptor beta subunit could only bind to Sbh2, but not Sbh1, in yeast microsomes [Jiang et al., 2008].

In contrast to Sbh2, which is degraded when not bound to the translocon, Sbh1 is stable on its own [Habeck et al., 2015]. Mutations of Sbh1 phosphorylation at T12, S20, and S35 to A, however, have an effect on its stability (Soromani et al., 2012). Differential regulation of the two specific signal peptide-binding Sbh subunits via channel-association or phosphorylation-induced stability changes, respectively, enables the yeast cell to independently control entry of specific subsets of proteins into the secretory pathway. This would allow for differential cell wall remodelling or secretion of specific enzymes in response to environmental signals by rapidly switching specificity of Sec61 and Ssh1 ER import channels. Metazoan Sec61β homologs, which often have longer IDRs and a larger number of phosphorylation sites [Kinch et al., 2002; Soromani et al., 2012], may have evolved to generate binding sites for specific subsets of signal peptides or Sec61 channel accessory factors by phosphorylation-controlled conformational switches to be able to rapidly adjust their secretory response to extracellular signals and physiological requirements [Bah et al., 2015].

In conclusion, our identification of the β subunit of the Sec61 channel as a specific signal peptide receptor has generated fundamental new insight into the function of this universally conserved entrance into the secretory pathway.

## EXPERIMENTAL PROCEDURES

### Molecular modelling of Sec61p-Sbh1 interaction

We postulated that the N-termini of Sec61p and Sbh1 may interact as a helix dimer and aimed at identifying putative contacts via molecular modelling. First, we modelled both the N-terminus of Sec61 (^1^MSSNRVLDLFKPFESFLPEVIAPE^24^) and the 36-aa N-terminal stretch of Sbh1 (^1^MSSPTPPGGQRTLQKRKQGSSQKVAASAPKKNTNSN^36^) into α-helical conformations using MODELLER [Webb and Sali, 2016]. Then, we integrated the N-terminus of Sec61p along with the downstream TMD1 into a POPC lipid bilayer using CHARMM-GUI [Jo et al., 2008] and the PPM server [Lomize et al., 2012] for orienting the proteins in the membrane. 150 mM KCl salt concentration was applied to make the overall system electrically neutral at pH 7.4. We performed a 300 ns long simulated tempering molecular dynamics (MD) simulation with the GROMACS 2018.8 package [van der Spoel et al., 2005] at 11 discrete temperatures spanning the range of 300 to 350 K to augment conformational sampling. The time-step was 2 fs, long-range electrostatic interactions were computed via PME, the force field was CHARMM36m with TIP3P water. Conformations with temperatures ≤ 303 K were extracted and clustered using a 2.5Å RMSD distance criterion. The representative conformation from the largest cluster was used for three subsequent plain MD simulations of 50 ns length each at 300 K in the presence of the N-terminal of Sbh1. We tested three putative contacting conformations between the cytosolic N-termini of Sbh1 and Sec61p having an angle between their helical axes of 0°, +30°, or -30°, respectively.

### Primers for site-directed mutagenesis

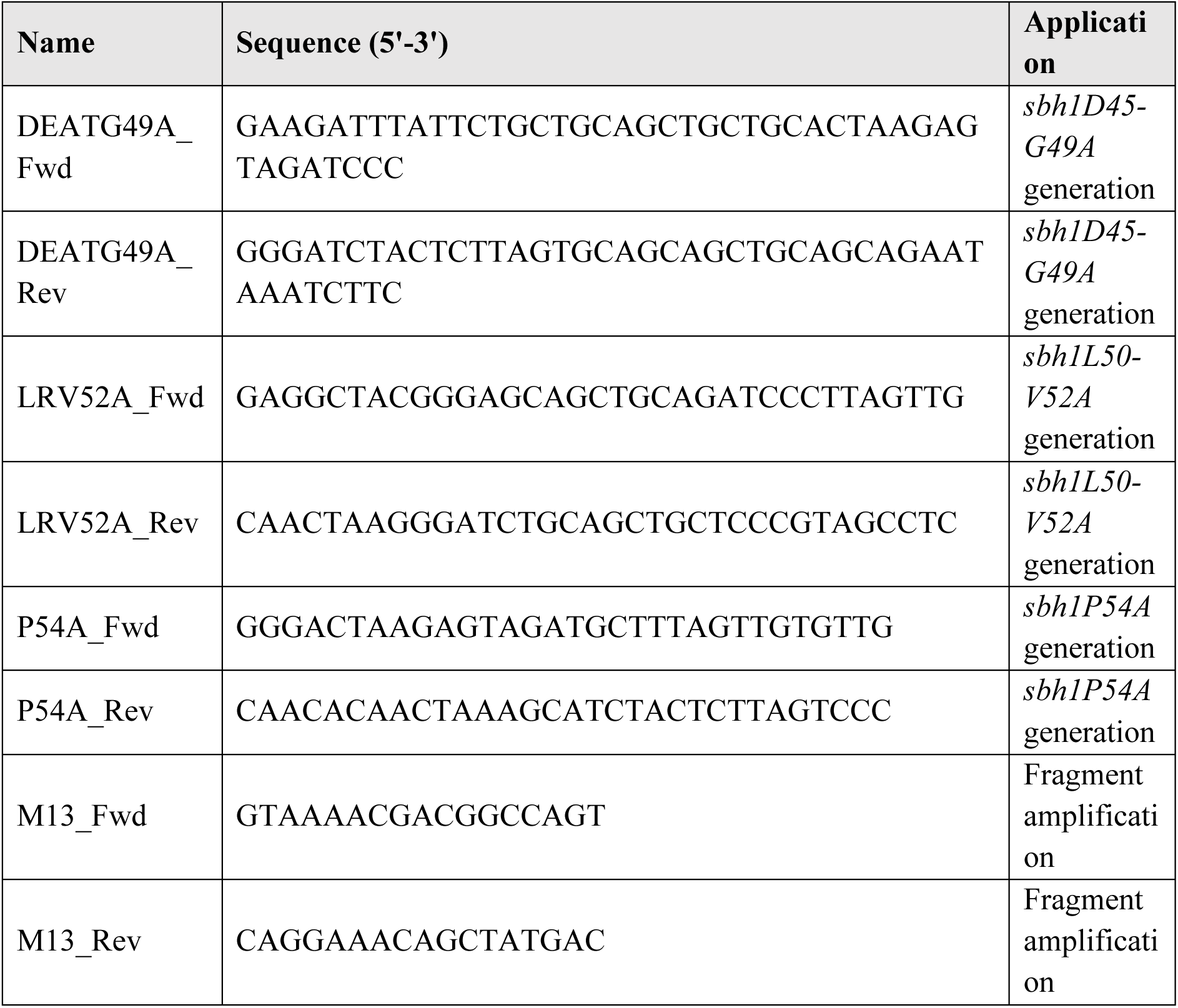

### Site-directed mutagenesis

The mutations in the membrane-proximal, conserved domain of Sbh1 were generated by site-directed mutagenesis of wildtype *SBH1* in pRS415 [Feng et al., 2007]. The *sbh1D45-G49A*, *sbh1L50-V52A,* and *sbh1P54A* mutants were made by substituting the respective amino acid residues with alanine. Forward and reverse primers for each mutation are listed below; they were first amplified separately with M13 primer in the opposite direction and *SBH1* coding sequence as template to generate “megaprimers”. These respective megaprimers were amplified together to generate the mutated *sbh1* insert fragments. The *sbh1D45-G49A* and *sbh1L50-V52A* fragments were cloned into pRS415 (CEN, *LEU2*) using ApaI and BamHI, and *sbh1P54A* was cloned using PstI and BamHI. Mutations were confirmed by DNA sequencing. Plasmids were transformed into KRY588 (*MATa sbh1::KanMx. sbh2::hphMx leu2-3,112 ura3-52 GAL+*) [Barbieri et al., 2023] and selected on synthetic complete medium without leucine at 30 °C.

### Yeast Growth Assays

All experiments with *sbh1* mutants were done in the KRY588 (*MATa sbh1::KanMx. sbh2::hphMx leu2-3,112 ura3-52 GAL+*) background transformed with the indicated SBH1 variants expressed from CEN plasmids [Barbieri et al., 2023]. The parental strain with *SBH1 SBH2* wildtype genes was used as control. An OD_600_ of 1 of yeast grown in selective medium to early exponential phase was harvested, washed with sterile deionised water and serially 10-fold diluted 4 times for a total of 5 concentrations. For each dilution, 5 µl (equivalent to 10^5^-10 cells) was dropped onto synthetic complete medium minus Leu agar plates and grown at 20 °C, 30 °C and 38 °C. Growth was documented after 3 days.

### Preparation and Analysis of Cell Extracts

For detection of translocation defects, cells were grown to an OD_600_ of 0.5 at 24 °C and shifted to 38 °C for 3 hours. Cells (2 OD_600_) were harvested at 600 x g for 1 min and supernatants were discarded. Pellets were washed with 1 ml of sterile deionized water, resuspended in 100 µl 2x SDS-PAGE sample buffer (100 mM Tris-HCl, pH 6.8 / 4 % SDS / 0.2 % bromophenol blue / 20 % glycerol / 200 mM DTT). Glass beads (50 µl by volume, acid washed, 1 mm, Sigma) were added and the cells were disrupted in a bead beater (Mini-BeadBeater-24, BioSpec Products Inc.) at 4 °C for 1 min twice, with 1 min pause in between cycles. Samples were denatured at 65 °C for 10 min for membrane proteins or at 95 °C for 5 min for soluble proteins, and centrifuged at 11,000 x g for 1 min. Samples were analysed in pre-cast 4-12% gradient Bis-Tris gels (NuPAGE Novex, Invitrogen), followed by Western Blotting using an appropriate concentration of primary antibody, followed by anti-rabbit-HRP secondary antibody (1:10,000, Sigma-Aldrich). Immunodetection was done using the SuperSignal West Pico PLUS chemiluminescent substrate (Thermo Fisher Scientific). Signals were detected using the AI600 Imager (Amersham) and quantified using ImageQuant™ TL software (GE Healthcare).

### Antibodies used in Western Blotting

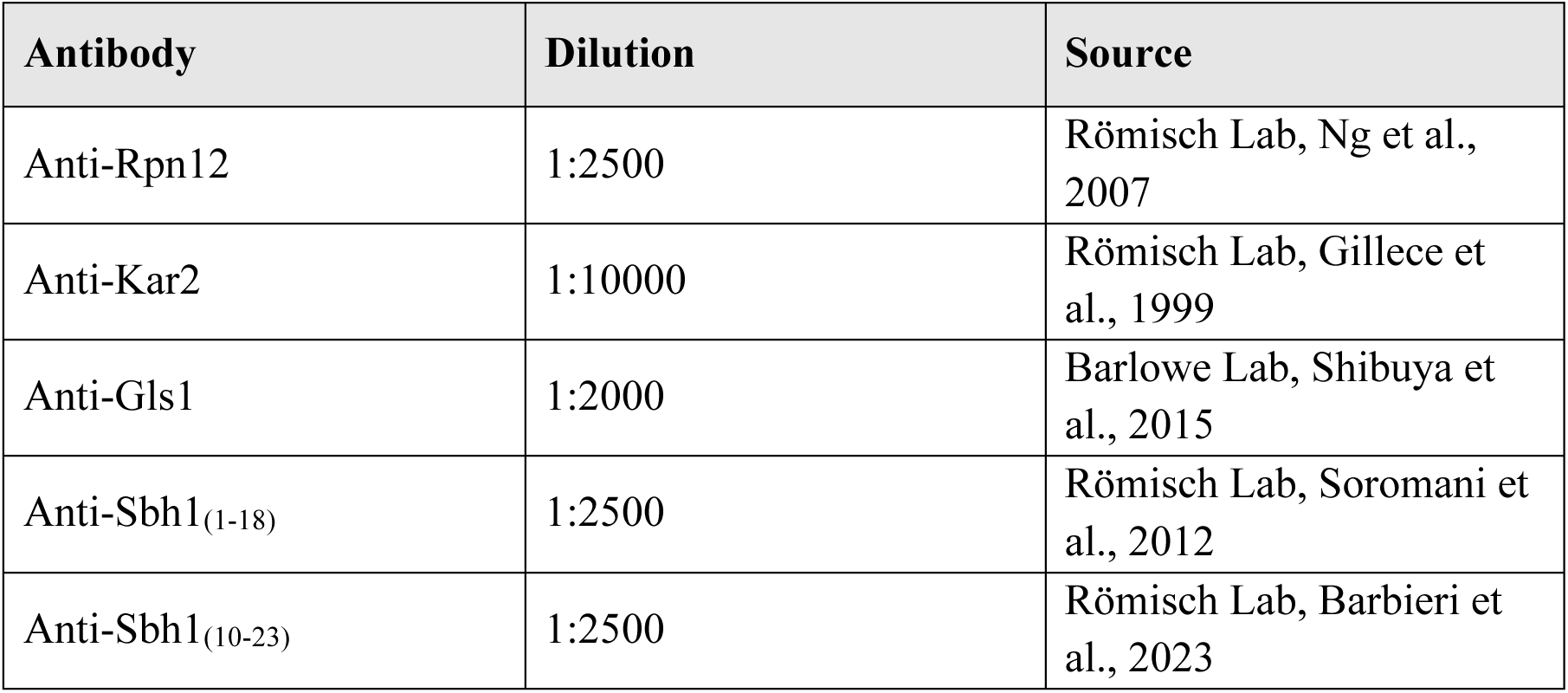
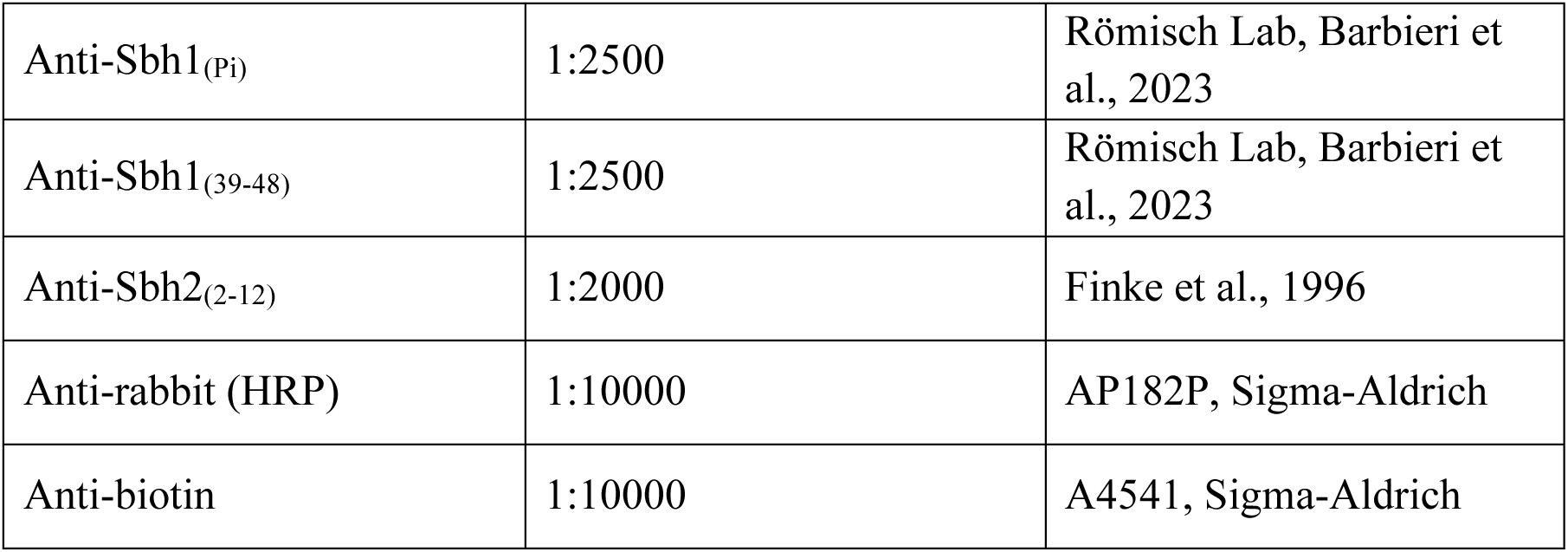

### Synthetic Sbh1 cytosolic domain phosphorylated variants

| Protein | Sequence |
| --- | --- |
| Sbh1 S3P | Ac-S(pS)PTPPGGQRTLQKRKQGSSQKVAASAPKKNTNSNNSI<br>LKI YSDEATGLRVDP |
| Sbh1 T5P | Ac-SSP(pT)PPGGQRTLQKRKQGSSQKVAASAPKKNTNSNNSI<br>LKI YSDEATGLRVDP |
| Sbh1 S3PT5P | Ac-S(pS)P(pT)PPGGQRTLQKRKQGSSQKVAASAPKKNTNSN<br>SIL KIYSDEATGLRVDP |

### Biotinylated synthetic signal peptides used for crosslinking

| Peptide | Sequence |
| --- | --- |
| Gls1 SP (1-24) | MLISKSKMFKTFWILTSIVLLASA-KBiotin |
| Irc22 SP (1-21) | MRFSMLIGFNLLTALSSFCAA-KBiotin |
| Carboxypeptidase Y (CPY) SP (1-20) | MKAFTSLLCGLGLSTTLAKA-KBiotin |
| prepro-alpha Factor (ppaF) SP (1-19) | MRFPSIFTAVLFAASSALA-KBiotin |
| Mpd2 SP (1-22) | MKLHGFLFSVLSTCVVILPALA-KBiotin |
| Pho3 SP (1-19) | MFKSVVYSVLAAALVNAGT-KBiotin |
| Pdi1 SP (1-22) | MKFSAGAVLSWSSLLLASSVFA-KBiotin |
| Sec71/1 TMD<br>(26-50) | KSISVYTPLIYVFILVVSLVMFASS-KBiotin |
| Sec71/2 TMD<br>(28-54) | ISVYTPLIYVFILVVSLVMFASSYRKK-KBiotin |

### Peptide Synthesis

Biotinylated signal peptides were synthesized with an automated ResPepSL synthesiser (Intavis) using amide Rink resin as the solid phase and Fmoc-protected amino acids (Carbolution) for coupling according to the method of Merrifield [Merrifield, 1963]. Fmoc-L-Lys-(Biotin) was used as starting amino acid at the C-terminal end. The resin-coupled peptides were cleaved and deprotected with trifluoroacetic acid (Sigma Aldrich) followed by precipitation with tert-butyl methyl ether (Fisher Scientific) and further analysis and purification by preparative HPLC (Merck) (purtity > 90%). Lyophilized peptides (Zirbus, VaCo5) were stored at 4 °C. Phosphorylated forms of the Sbh1 cytosolic domain were synthesized commercially by Peptide Specialty Laboratories, Heidelberg.

### Pulse-Labelling and Immunoprecipitation

Pulse labeling protocol was adapted from Barbieri *et al*., 2023. Yeast strains were grown in YPD or selective synthetic complete media at their permissive temperature (24 °C or 30 °C) to an OD_600_ of 0.5 and transferred to their respective restrictive temperature (38 °C or 20 °C) for 3 hours. Cells were harvested at 4500 g for 5 mins, washed twice with Labelling Medium (0.67% Yeast Nitrogen Base without amino acids and ammonium sulfate / 5% Glucose / auxotrophy supplements as required) and resuspended to OD_600_ of 6/ml using Labelling Medium. Aliquots of 1.5 OD_600_ of each strain were pre-incubated at their restrictive temperatures for 10 min at 800 rpm in a Thermoblock (Eppendorf). Proteins were labelled for 5 mins (for soluble proteins) or 15 mins (for membrane proteins) with 30 µCi of Express Protein Labelling Mix (Perkin Elmer) per sample at their restrictive temperature. The pulse was stopped by killing the cells using cold Tris-Azide Buffer (20 mM Tris-HCl, pH 7.5 / 20 mM NaN_3_) and immediate transfer to ice. Cells were harvested for 1 min, 4 °C, at full speed in an Eppendorf 5424-R microfuge, washed once with Tris-Azide buffer, and resuspended in Resuspension Buffer (100 mM Tris-HCl, pH 9.4 / 10 mM DTT/ 20 mM NaN_3_), incubated for 10 mins at room temperature, and collected at full speed for 1 min at 4 °C. Cells were resuspended in Lysis Buffer (20 mM Tris-HCl, pH 7.5 / 0.5% SDS / 1 mM DTT / 1 mM PMSF). An equal volume of glass beads was added to each sample, and cells were disrupted by bead-beating twice for 1 min, with 1 min interval at room temperature. Samples were denatured at 90 °C (for soluble proteins) or 65 °C (membrane proteins). The beads were washed thrice with 250 µL Washing Buffer (150 mM NaCl / 1% Triton X-100 / 15 mM Tris-HCl, pH 7.5 / 2 mM NaN_3_ / 1 mM PMSF). The supernatant for each sample was collected from each wash, combined, and subjected to immunoprecipitation.

Non-specifically binding proteins were cleared from each sample by incubation with 60 µL of 20% Protein A-Sepharose™ CL-4B (GE Healthcare) in IP Buffer (150 mM NaCl / 1% Triton X-100 / 15 mM Tris-HCl, pH 7.5 / 2 mM NaN_3_ / 0.1% SDS) and incubation for 30 min under rotation at room temperature. Supernatants were collected by a quick spin in an Eppendorf 5424-R microfuge and incubated overnight at 4 °C under rotation with 20% Protein A-Sepharose™ CL-4B in IP Buffer and 10 µl of anti-CPY (Römisch lab) or anti-DPAPB (Martin Pool, Manchester) polyclonal rabbit antisera. The beads were collected by a quick spin and washed twice with IP Buffer, twice with Urea Wash (2 M Urea / 200 mM NaCl / 1 % Triton X-100 / 100 mM Tris-HCl, pH 7.5 / 2 mM NaN_3_), once with Con A Wash (500 mM NaCl / 1 % Triton X-100 / 20 mM Tris-HCl, pH 7.5 / 2 mM NaN_3_) and once with Tris-NaCl Wash (50 mM NaCl / 10 mM Tris-HCl, pH 7.5 / 2 mM NaN_3_). Beads were collected by a quick spin and resuspended in 2x SDS-PAGE sample buffer. Samples were denatured at 95 °C (for soluble proteins) or 65 °C (membrane proteins), loaded onto 4-12% pre-cast Bis-Tris gels and subjected to electrophoresis. Gels were fixed for 30 mins in a Fixing Solution (10% Acetic acid / 40% Methanol), washed with deionized water, and dried for 1 hour at 80 °C in a gel dryer (Model 583, Bio-Rad). The dried gels were exposed to phosphorimaging plates for 72 hours. The signal was acquired using Typhoon Trio™ Variable Mode Imager (GE Healthcare) and analyzed in ImageQuant™ TL software.

### Sbh1/Sbh2 cytosolic domain expression and purification

DNA encoding the cytosolic domains of Sbh1, Sbh1 cysteine variants, and Sbh2 was codon-optimized for expression in *E. coli*, cloned into pET24a, and expressed in *E. coli* BL21(DE3)RIL. A 2.5 L culture was grown in TB-FB medium supplemented with 1.5% lactose, 0.05% glucose, 2 mM MgSO4, 30 µg/ml kanamycin, and 35 µg/ml chloramphenicol at 37 °C until an OD_600_ of 0.8 was reached, then shifted to 18 °C and incubated overnight. Cells were harvested, washed once in Ni running buffer (50 mM Tris-HCl, pH 8.0, 500 mM NaCl, 30 mM imidazole, 5 mM β-mercaptoethanol), and resuspended in 30 ml of the same buffer. The suspension was lysed by sonication on ice for 5 min using a Branson Ultrasonics Sonifier Model 450, and the lysate was cleared by centrifugation at 48,000 × g for 30 min at 4 °C. The supernatant was loaded onto a HisTrap Crude column (Cytiva) using an ÄKTA Prime chromatography system (Cytiva) at a flow rate of 1 ml/min. After washing with Ni running buffer until a stable baseline was reached, the protein was eluted with a step gradient of 100% Ni elution buffer (50 mM Tris-HCl, pH 8.0, 500 mM NaCl, 350 mM imidazole, 5 mM β-mercaptoethanol). The eluate containing Sbh protein was concentrated to 1 ml using an Amicon Ultra centrifugal filter (MWCO 3 kDa, Sigma-Aldrich), adjusted to a final volume of 10 ml with Ni running buffer, and supplemented with 0.5 mg 10xHis-SuperTEV (https://dx.doi.org/10.17504/protocols.io.yxmvm2zxng3p/v1). Samples were cleaved overnight at 4 °C. To remove non-cleaved protein and protease, the samples were applied to a second HisTrap Crude column, and the flow-through was collected. For size-exclusion chromatography, the sample was concentrated to 5 ml, loaded onto a HiLoad Superdex 75 16/60 column (Cytiva), and eluted in SEC buffer (20 mM HEPES-KOH, pH 6.8, 150 mM potassium acetate, 5 mM magnesium acetate, 250 mM sorbitol, and 2 mM TCEP) at 0.5 ml/min. The eluate was adjusted to 1 mg/ml protein and stored in aliquots at -80 °C.

### Sbh1/Sbh2 cytosolic domain crosslinking to signal peptides

Disuccinimidyl suberate (DSS) and bismaleimidohexane (BMH), as well as biotinylated peptides, were dissolved in DMSO to 15 mM. Crosslinking reactions (100 µl) contained 15 µM Sbh protein, 150 µM signal peptide, 2% DMSO, 20 mM HEPES-KOH, pH 7.4, 150 mM potassium acetate, 5 mM magnesium acetate, and 250 mM sorbitol. Reactions were initiated by addition of DSS or BMH to 150 µM final concentration and incubated for 30 min at 20 °C. Crosslinking was quenched with 50 µl 1 M Tris-HCl, pH 7.5, for 15 min at room temperature. Samples were mixed with an equal volume of 2× sample buffer, heated at 95 °C for 5 min, and separated by Tricine-SDS-PAGE as described previously Schägger, 2006. Proteins were transferred to nitrocellulose membranes and probed as described above. For mass spectrometry, reactions were prepared identically, except that concentrations of Sbh protein, signal peptide, and crosslinker were increased fourfold.

### Preparation and phosphatase treatment of crude yeast microsomes

Crude microsomes were prepared and desphosphorylated as described in Barbieri et al., 2023.

### Signal peptide crosslinking to Sbh1 in purified microsomes

Sucrose gradient-purifed microsomes were prepared as in Pilon et al. (1997). Prior to crosslinking, peripheral proteins including ribosomes and signal recognition particles were stripped off the membranes by successive incubations on ice with 50 mM EDTA buffer, followed by 500 mM potassium acetate buffer as described in Ng et al. (1996). Subsequently, 80 µl membranes (OD_280_ = 30/ml) were treated with Lambda protein phosphatase (LambdaPP; New England Biolabs) or mock-treated according to the manufacturer’s instructions. After treatment membranes were harvested by centrifugation at 20,000 g for 20 min at 4 °C and resuspended in 80 µl crosslinking buffer (20 mM HEPES-KOH, pH 7.4, 150 mM potassium acetate, 5 mM magnesium acetate, 250 mM sorbitol). For crosslinking reactions 20 µl membranes were combined with crosslinking buffer and synthetic peptides (1.5 mM final concentration) and incubated at 20 °C for 20 min. Crosslinking reactions were started by the addition of DSS (1.5 mM final concentration) and incubated at 20 °C for 30 min. Reactions were terminated by the addition 8.4 M ammonium acetate to a final concentration of 600 mM. Samples were harvested by centrifugation, resuspended in 80 µl Tricine sample buffer, separated on Tricine gels, and transferred to nitrocellulose. Sbh1 and crosslinking products were detected with Sbh1 (39-48) antibody.

### Mass spectrometry analysis of crosslinked samples (XL-MS)

Cross-linked samples were lyophilized and resuspended in 50 μL of 8 M urea, reduced, alkylated, and digested with trypsin (Promega), as described previously [Leitner et al., 2014]. Digested peptides were purified by solid-phase extraction (50-mg SepPak cartridges, Waters), and cross-linked peptides were enriched by size exclusion chromatography using an ÄKTAmicro chromatography system (GE Healthcare) equipped with a SuperdexTM Peptide 3.2/30 column (column volume = 2.4 ml). For each cross-linked sample, four fractions (0.8-1.0, 1.0-1.1, 1.1-1.2, 1.2-1.4 ml) were collected. Absorption levels at 215 nm of each fraction were used to normalize peptide concentration prior to analysis by liquid chromatography-tandem mass spectrometry (LC-MS/MS) on an Orbitrap Eclipse Tribrid mass spectrometer (Thermo Electron, San Jose, CA). Peptides were loaded onto a C18 column (Acclaim™ PepMap™ AcclaimTM PepMapTM 100 C18-LC 50mm, Thermo Scientific) and separated on a Vanquish neo UHPLC system (Thermo Scientific) at a flow rate of 300 nL/min over a 48 min gradient optimized for each fraction (3 % to 6-15 % for 4 min, 6-15 % to 25-40 % in 35 min, 25-40 % to 35-45 % in 10 min and 35-45 % to 80 % acetonitrile in 0.1 % formic acid in 1 min). Full scan mass spectra were acquired in the Orbitrap at a resolution of 120,000, a scan range of 400–1500 *m/z*, and a maximum injection time of 50 ms. Most intense precursor ions (intensity ≥5.0 × 10^3^) with charge states 2–8 and monoisotopic peak determination set to “peptide” were selected for MS/MS fragmentation by CID at 35 % collision energy in a data-dependent mode. The duration for dynamic exclusion was set to 45 s. MS/MS spectra were analyzed in the Ion Trap at a rapid scan rate and maximum injection time in dynamic mode. For the crosslink analysis, a database was compiled which contained the amino acid sequence of Shb1 and respective peptide Msn1, Irc22 or Gls1. The data was analyzed using XQuest/xProphet pipeline^1^ allowing a maximum number of 2 miss cleavages. All unique cross- and mono-links with an ld-score > 20 and an FDR < 0.05 were considered for evaluation and links were visualized via xiNET [Combe et al., 2015].

## ACKNOWLEDGMENTS

Antibodies against DPAPB were kindly provided by Martin Pool (University of Manchester, UK), antisera against Gls1 were a gift from Charlie Barlowe (Dartmouth University, USA), Sbh2 antibody was a gift from Enno Hartmann (Lübeck University). We thank Aline Leguede and Yasaman Farahani for expert technical assistance and members of the Römisch lab for comments on the manuscript.

## SUPPLEMENTARY FIGURE LEGENDS

**Supplementary Figure 1.**
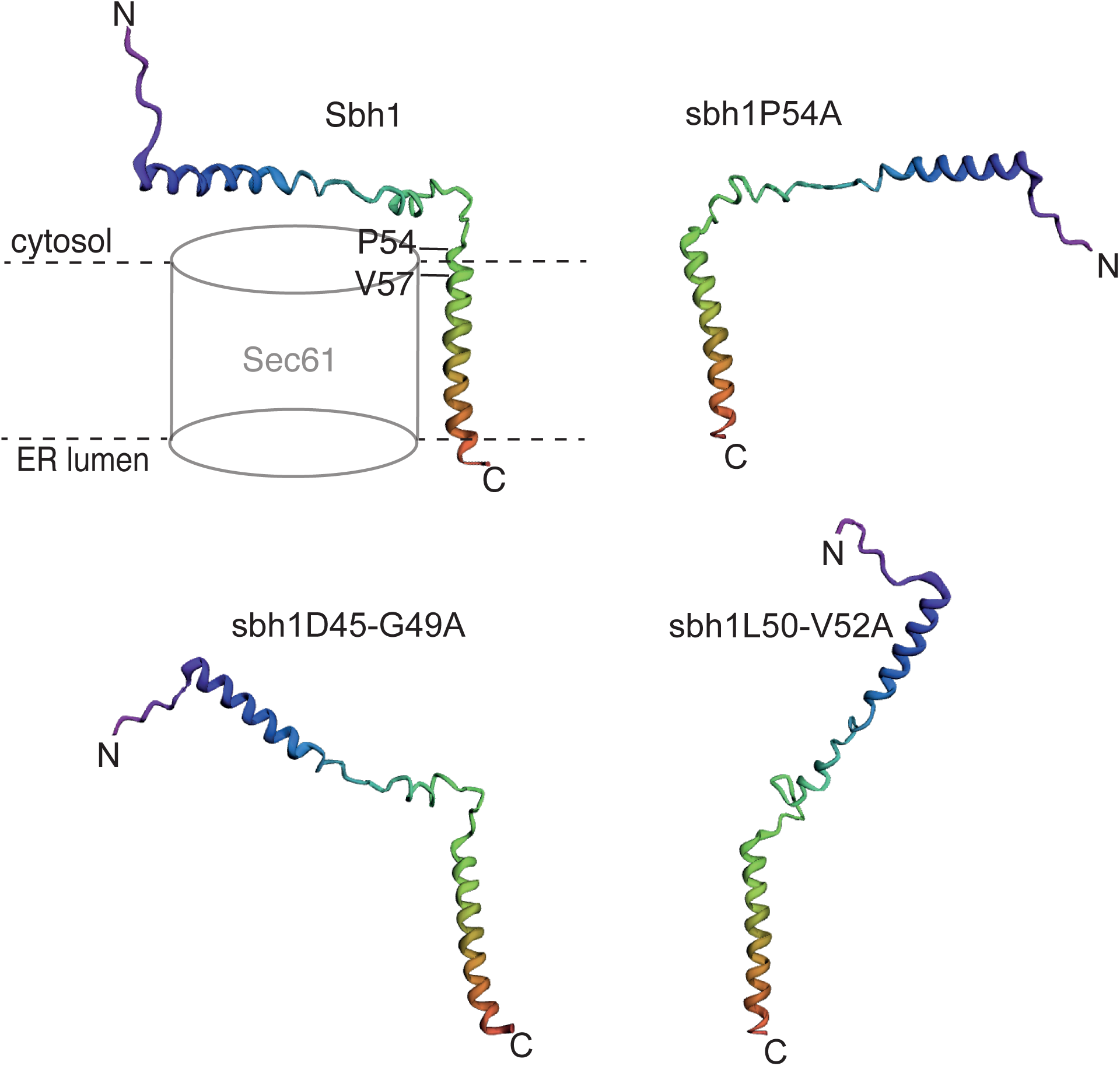
Conformations of Sbh1 wildtype and cytosolic domain mutant proteins. Sbh1 wildtype and sbh1P54A, sbh1L50-V52A, and sbh1D45-G49A mutants were subjected to Rosetta *ab initio* modelling [Das et al., 2008; Alford et al., 2017]. The resulting structures are shown with the C-terminal transmembrane helix in the same fixed position, such that changes in the orientation of the cytosolic domain due to the mutations are evident. Coloring is solely to highlight N- and C-termini, and does not indicate prediction confidence levels. The Sbh1 wildtype structure is the same as the one shown in the model in Fig. 4E.

**Supplementary Figure 2.**
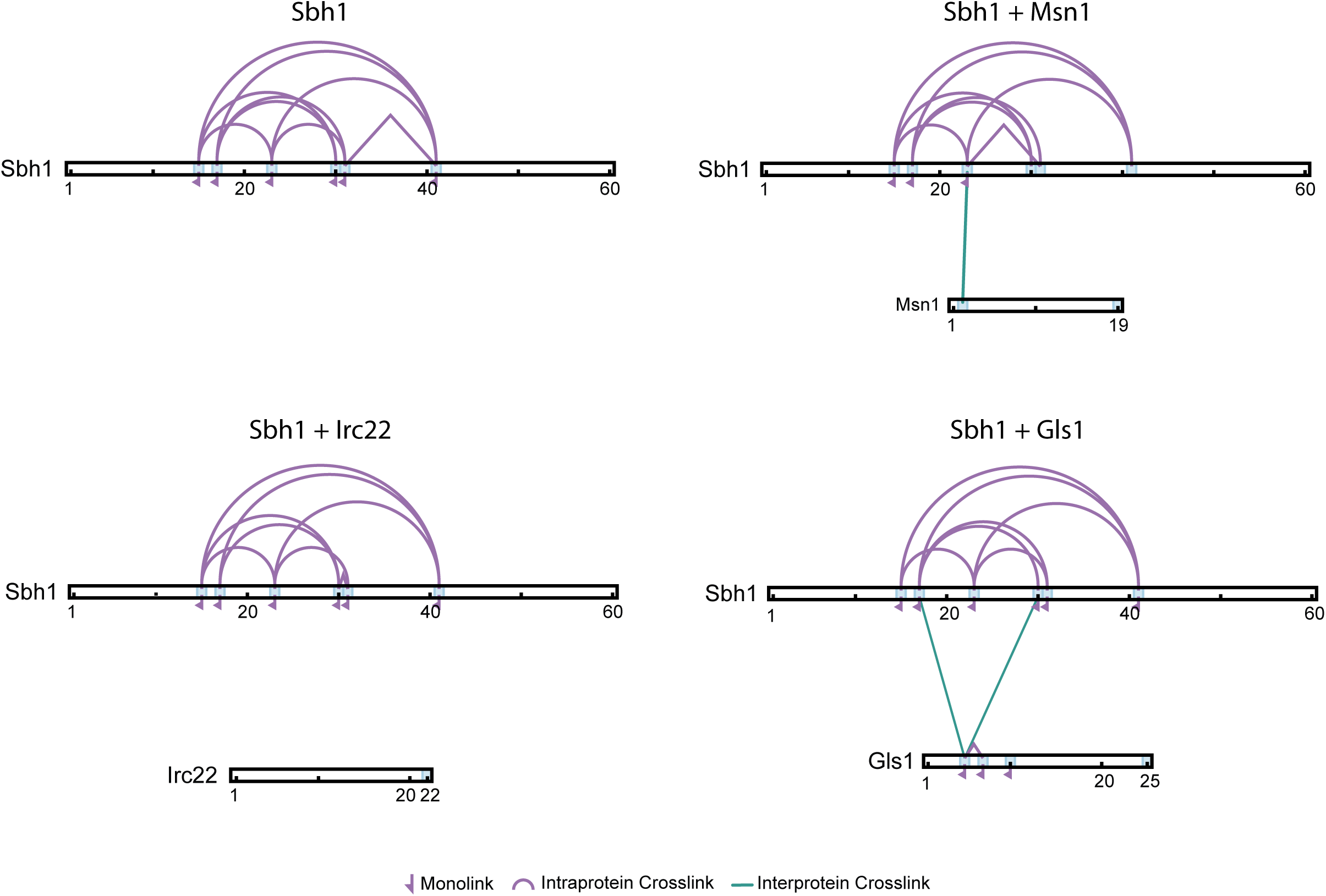
Crosslinking/mass spectrometry (XL-MS) analysis of Sbh1 cytosolic domain/signal peptide complexes. Sbh1 on its own or in the presence of the indicated signal peptides was incubated with amino-specific homobifunctional crosslinker DSS as described in the methods section. High-confidence crosslinks identified by XL-MS were mapped onto the sequence of the Sbh1 cytosolic domain and the sequences of Irc22, Mns1, and Gls1 signal peptides. Lysine residues are indicated by blue tick marks. Link types are defined within the figure.

**Supplementary Figure 3.**
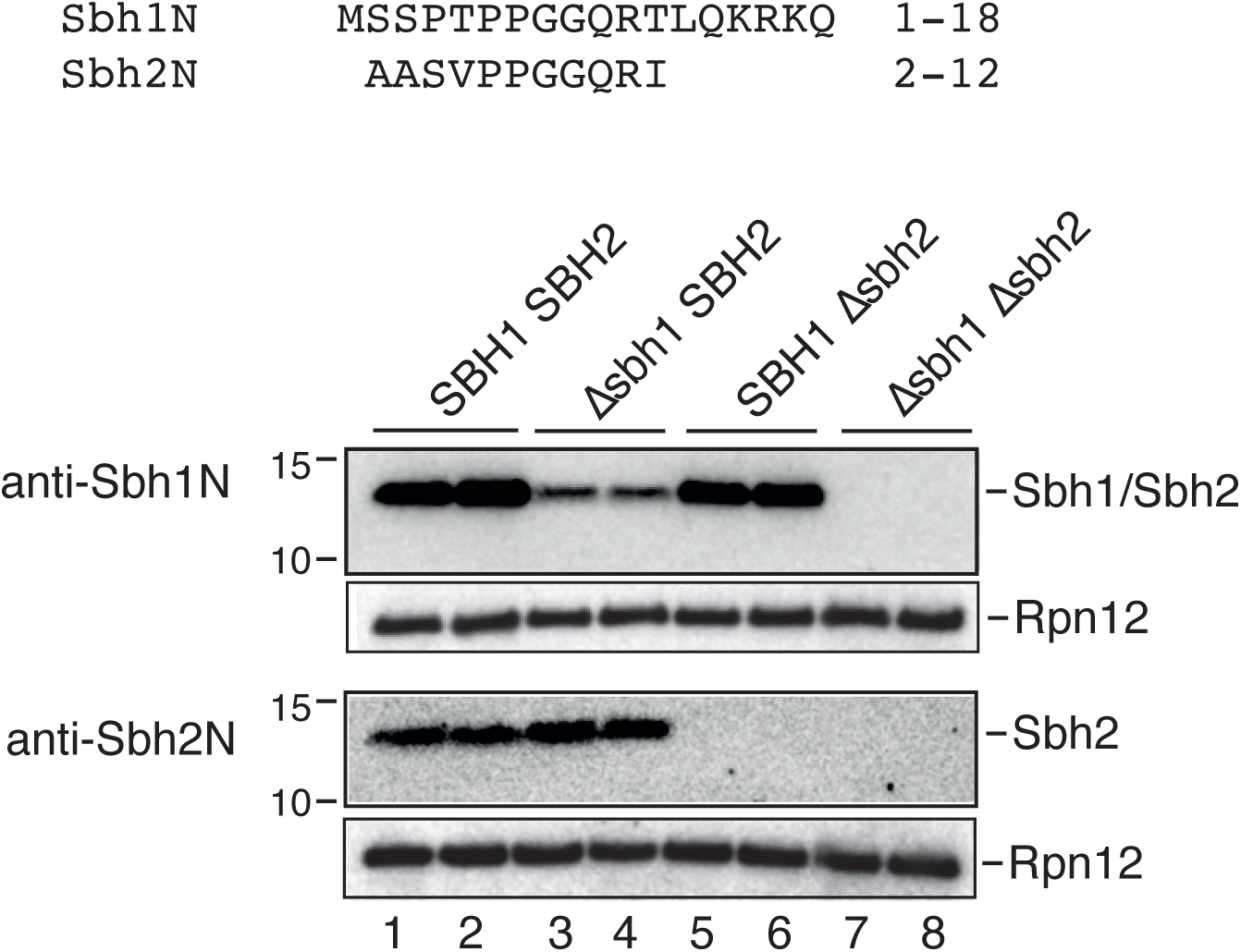
N-terminal antibody reactivity against Sbh1 and Sbh2. Antibodies were raised against the synthetic peptides shown on top of the figure, which are derived from the N-termini of Sbh1 and Sbh2, respectively, coupled via an additional C residues at the C-termini. Equal amounts of extracts of the indicated strains were separated on 4-20 % SDS gradient gels, transferred to nitrocellulose and detected with the indicated antibodies. Rpn12 was used as a loading control.

## SUPPLEMENTARY TABLES

**Supplementary Table 1. Cross-linking mass spectrometry (XL-MS) analysis of SP/Sbh1 complexes.**

