## Supplementary figures and images for "Sbh1/Sec61β is a specific signal peptide receptor within the Sec61 channel"

### Supp Fig 1

#### Supplementary Figure 1

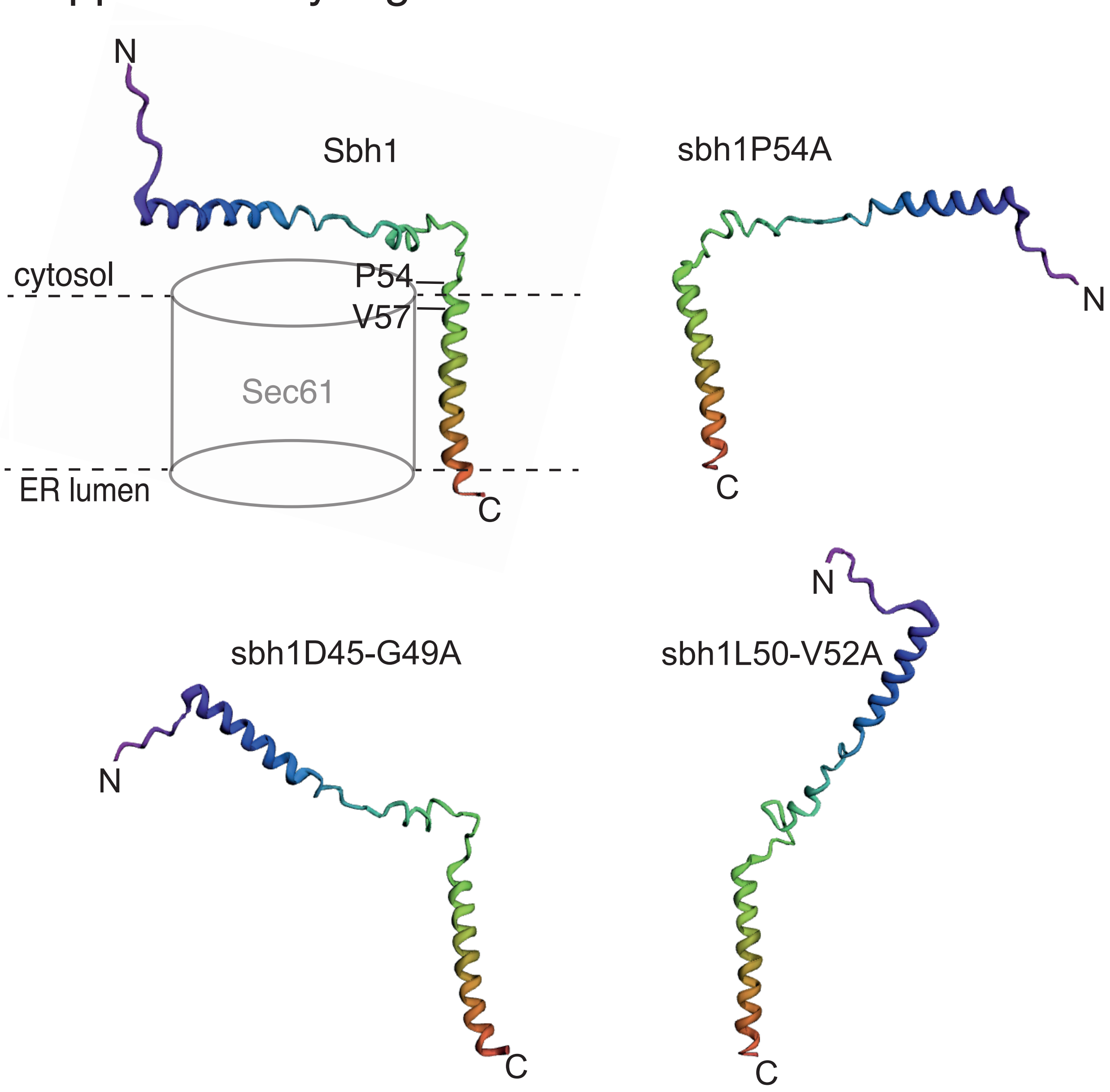

### Supp Fig 2

#### Supplementary Figure 2

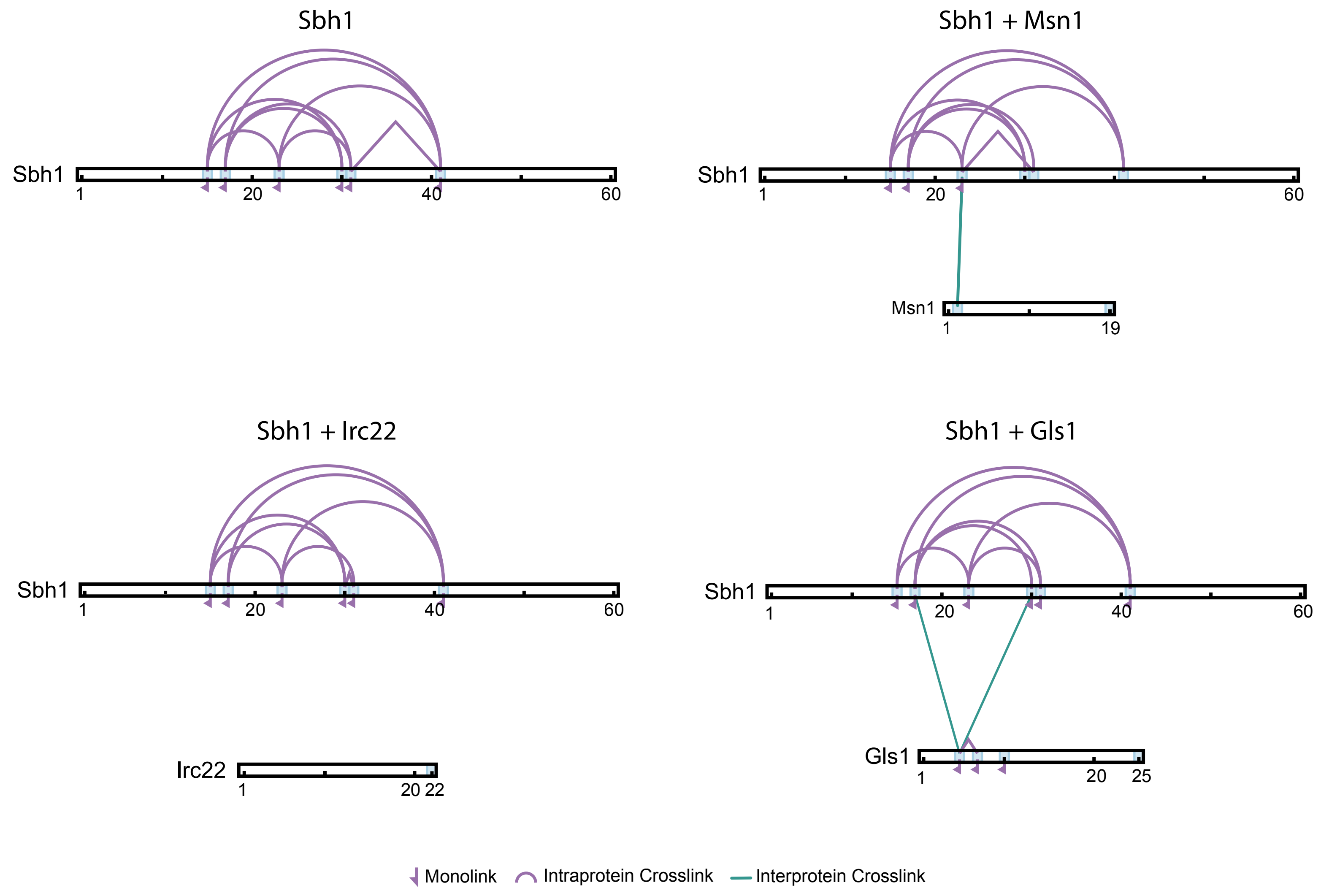

### Supp Fig 3

#### Supplementary Figure 3

```
Sbh1N     MSSPTPPGGQRTLQKRKQ  1-18
Sbh2N      AASVPPGGQRI        2-12
```

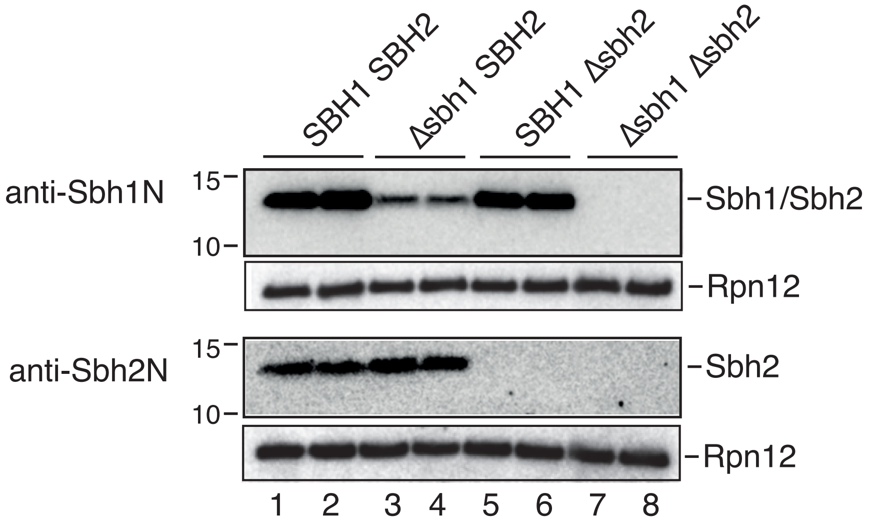
