## Supplementary material for "Sbh1/Sec61β is a specific signal peptide receptor within the Sec61 channel": Supp Table 1

***Legend***

| | |
|---|---|
| *Crosslinked Peptide* | amino acid sequence of peptide a and peptide b of crosslinked peptide including the position of crosslinking sites in peptide a and peptide b |
| *Protein1* | name of protein to which peptide a belongs |
| *Protein2* | name of protein to which peptide b belongs |
| *XLType* | type of linker attachment (inter-protein crosslink, intra-protein crosslink, intralink or monolink) |
| *Pos1* | absolute position of lysine residue in peptide a of protein 1 which was crosslinked (absolute position without affinity tag used for purification) |
| *Pos2* | absolute position of lysine residue in peptide b of protein 2 which was crosslinked (absolute position without affinity tag used for purification) |
| *Mr* | precursor ion mass |
| *Mz* | m/z of multiply charged precursor ion |
| *z* | charge state of identified crosslinked peptide |
| *deltaS* | delta score is a measure for how close the best assigned hit scored in regard to second best |
| *ld-Score* | linear discriminant score; scoring scheme to discriminate between true positive crosslinks and false positive hits |
| *Biological Replicate* | indicates in which biological replicate (I, II, III) the peptide was identified |

| *Crosslinked Peptide* | *Protein1* |
|---|---|
| KQGSSQKVAASAPK-K7-155 | Sbh1 |
| KQGSSQKVAASAPK-K1-K7 | Sbh1 |
| KQGSSQKVAASAPK-K7-156 | Sbh1 |
| KQGSSQKVAASAPK-KQGSSQK-a7-b1 | Sbh1 |
| VAASAPKK-K7-156 | Sbh1 |
| NTNSNNSILKIYSDEATGLR-K10-156 | Sbh1 |
| VAASAPKK-K8-156 | Sbh1 |
| KQGSSQKVAASAPK-KNTNSNNSILK-a7-b1 | Sbh1 |
| NTNSNNSILKIYSDEATGLR-K10-155 | Sbh1 |
| KNTNSNNSILKIYSDEATGLR-K11-155 | Sbh1 |
| VAASAPKK-K7-155 | Sbh1 |
| KNTNSNNSILKIYSDEATGLR-K1-K11 | Sbh1 |
| NTNSNNSILKIYSDEATGLR-VAASAPKK-a10-b7 | Sbh1 |
| VAASAPKK-K8-155 | Sbh1 |
| KNTNSNNSILKIYSDEATGLR-QGSSQKVAASAPK-a11-b6 | Sbh1 |
| QGSSQKVAASAPK-TLQKRK-a6-b4 | Sbh1 |
| KNTNSNNSILKIYSDEATGLR-K11-156 | Sbh1 |
| QGSSQKVAASAPK-K6-156 | Sbh1 |
| KNTNSNNSILK-K1-156 | Sbh1 |
| VAASAPKK-K7-K8 | Sbh1 |
| NTNSNNSILKIYSDEATGLRVDPENLYFQ-K10-156 | Sbh1 |
| RKQGSSQK-K2-156 | Sbh1 |
| KNTNSNNSILK-K1-155 | Sbh1 |
| VAASAPKKNTNSNNSILK-K7-K8 | Sbh1 |
| TLQKRK-K4-156 | Sbh1 |
| KQGSSQKVAASAPK-QGSSQKVAASAPK-a7-b6 | Sbh1 |
| KNTNSNNSILKIYSDEATGLR-KQGSSQKVAASAPK-a11-b7 | Sbh1 |
| KNTNSNNSILK-K11-156 | Sbh1 |
| NTNSNNSILKIYSDEATGLR-TLQKRK-a10-b4 | Sbh1 |
| NTNSNNSILKIYSDEATGLR-KQGSSQK-a10-b1 | Sbh1 |
| KQGSSQK-K1-155 | Sbh1 |
| KQGSSQK-K1-156 | Sbh1 |

| | |
|---|---|
| QGSSQKVAASAPK-K6-155 | Sbh1 |
| VAASAPKK-KQGSSQK-a7-b1 | Sbh1 |
| KNTNSNNSILKIYSDEATGLR-KQGSSQK-a11-b1 | Sbh1 |
| NTNSNNSILKIYSDEATGLR-RKQGSSQK-a10-b2 | Sbh1 |
| QGSSQKVAASAPK-KQGSSQK-a6-b1 | Sbh1 |
| KQGSSQKVAASAPK-VAASAPKK-a7-b7 | Sbh1 |
| KQGSSQKVAASAPK-K7-K14 | Sbh1 |
| RKQGSSQK-VAASAPKK-a2-b7 | Sbh1 |
| QGSSQKVAASAPKK-K6-156 | Sbh1 |
| KQGSSQKVAASAPK-K1-156 | Sbh1 |
| QGSSQKVAASAPKK-TLQKRK-a6-b4 | Sbh1 |
| KNTNSNNSILKIYSDEATGLR-K1-156 | Sbh1 |
| VAASAPKKNTNSNNSILK-RKQGSSQK-a7-b2 | Sbh1 |
| QGSSQKVAASAPK-KNTNSNNSILK-a6-b1 | Sbh1 |
| KNTNSNNSILK-KQGSSQK-a1-b1 | Sbh1 |
| KQGSSQK-TLQKRK-a1-b4 | Sbh1 |
| VAASAPKK-TLQKRK-a7-b4 | Sbh1 |
| NTNSNNSILKIYSDEATGLRVDPENLYFQ-K10-155 | Sbh1 |
| QGSSQKVAASAPK-K6-K13 | Sbh1 |
| QGSSQKVAASAPKK-K6-K13 | Sbh1 |
| KNTNSNNSILK-RKQGSSQK-a1-b2 | Sbh1 |
| VAASAPKKNTNSNNSILK-K7-156 | Sbh1 |
| KNTNSNNSILK-VAASAPKK-a1-b7 | Sbh1 |
| KNTNSNNSILKIYSDEATGLR-QGSSQKVAASAPK-a1-b6 | Sbh1 |
| VAASAPKKNTNSNNSILK-K8-156 | Sbh1 |
| NTNSNNSILKIYSDEATGLR-QGSSQKVAASAPK-a10-b6 | Sbh1 |
| KNTNSNNSILKIYSDEATGLR-VAASAPKK-a11-b7 | Sbh1 |
| VAASAPKKNTNSNNSILK-K7-K18 | Sbh1 |

| *Protein2* | *XLType* | *Pos1* | *Pos2* | *Mr* | *Mz* | *z* | *deltaS* | *ld-Score* |
|---|---|---|---|---|---|---|---|---|
| - | monolink | 23 | n/a | 1540,855 | 771,435 | 2 | 0,44 | 54,56 |
| - | intralink | 17 | 23 | 1523,827 | 508,95 | 3 | 0,33 | 53,64 |
| - | monolink | 23 | n/a | 1541,839 | 771,927 | 2 | 0,4 | 53,41 |
| Sbh1 | intra-protein › | 23 | 17 | 2285,234 | 572,316 | 4 | 0,93 | 48,62 |
| - | monolink | 30 | n/a | 926,543 | 464,279 | 2 | 0,65 | 47,6 |
| - | monolink | 41 | n/a | 2365,177 | 789,4 | 3 | 0 | 46,98 |
| - | monolink | 31 | n/a | 926,543 | 464,28 | 2 | 0,62 | 46,43 |
| Sbh1 | intra-protein › | 23 | 31 | 2755,481 | 552,104 | 5 | 0,66 | 46,33 |
| - | monolink | 41 | n/a | 2364,191 | 789,072 | 3 | 0 | 45,98 |
| - | monolink | 41 | n/a | 2492,283 | 1247,15 | 2 | 0 | 45,86 |
| - | monolink | 30 | n/a | 925,559 | 463,787 | 2 | 0,5 | 45,11 |
| - | intralink | 31 | 41 | 2475,257 | 826,093 | 3 | 0 | 44,98 |
| Sbh1 | intra-protein › | 41 | 30 | 3117,632 | 780,416 | 4 | 0,79 | 44,27 |
| - | monolink | 31 | n/a | 925,559 | 463,787 | 2 | 0,59 | 44,23 |
| Sbh1 | intra-protein › | 41 | 23 | 3732,926 | 747,593 | 5 | 0,87 | 43,51 |
| Sbh1 | intra-protein › | 23 | 15 | 2168,229 | 434,654 | 5 | 0 | 43,49 |
| - | monolink | 41 | n/a | 2493,272 | 1247,644 | 2 | 0 | 43,4 |
| - | monolink | 23 | n/a | 1413,748 | 707,882 | 2 | 0 | 43,33 |
| - | monolink | 31 | n/a | 1387,73 | 694,873 | 2 | 0 | 43,14 |
| - | intralink | 30 | 31 | 908,532 | 455,274 | 2 | 0,65 | 42,19 |
| - | monolink | 41 | n/a | 3470,684 | 1157,902 | 3 | 0 | 42,11 |
| - | monolink | 17 | n/a | 1073,583 | 537,799 | 2 | 0,86 | 42 |
| - | monolink | 31 | n/a | 1386,745 | 694,38 | 2 | 0 | 41,87 |
| - | intralink | 30 | 31 | 1994,081 | 665,701 | 3 | 0,13 | 41,28 |
| - | monolink | 15 | n/a | 928,569 | 465,293 | 2 | 0 | 41,11 |
| Sbh1 | intra-protein › | 23 | 23 | 2781,496 | 696,382 | 4 | 0,79 | 41,06 |
| Sbh1 | intra-protein › | 41 | 23 | 3861,028 | 1288,017 | 3 | 0,73 | 40,37 |
| - | monolink | 41 | n/a | 1387,73 | 694,873 | 2 | 0 | 40,34 |
| Sbh1 | intra-protein › | 41 | 15 | 3119,655 | 624,939 | 5 | 0,24 | 40,08 |
| Sbh1 | intra-protein › | 41 | 17 | 3108,569 | 778,15 | 4 | 0,88 | 39,46 |
| - | monolink | 17 | n/a | 916,497 | 459,257 | 2 | 0,69 | 39,36 |
| - | monolink | 17 | n/a | 917,481 | 459,748 | 2 | 0,73 | 39,1 |

| | | | | | | | | |
|---|---|---|---|---|---|---|---|---|
| - | monolink | 23 | n/a | 1412,761 | 707,388 | 2 | 0 | 38,88 |
| Sbh1 | intra-protein › | 30 | 17 | 1669,936 | 557,653 | 3 | 0,93 | 38,34 |
| Sbh1 | intra-protein › | 41 | 17 | 3236,665 | 1079,896 | 3 | 0,93 | 37,8 |
| Sbh1 | intra-protein › | 41 | 17 | 3264,671 | 1089,231 | 3 | 0 | 37,69 |
| Sbh1 | intra-protein › | 23 | 17 | 2157,14 | 540,293 | 4 | 0,94 | 37,53 |
| Sbh1 | intra-protein › | 23 | 30 | 2294,296 | 765,773 | 3 | 0,88 | 37,02 |
| - | intralink | 23 | 30 | 1523,83 | 762,923 | 2 | 0,67 | 37,01 |
| Sbh1 | intra-protein › | 17 | 30 | 1826,035 | 609,686 | 3 | 0,85 | 36,93 |
| - | monolink | 23 | n/a | 1541,84 | 771,928 | 2 | 0,9 | 36,88 |
| - | monolink | 17 | n/a | 1541,843 | 771,929 | 2 | 0,12 | 36,58 |
| Sbh1 | intra-protein › | 23 | 15 | 2296,321 | 460,272 | 5 | 0,67 | 36,09 |
| - | monolink | 31 | n/a | 2493,28 | 1247,648 | 2 | 0 | 35,52 |
| Sbh1 | intra-protein › | 30 | 17 | 2911,58 | 971,534 | 3 | 0,35 | 35,14 |
| Sbh1 | intra-protein › | 23 | 31 | 2627,39 | 657,855 | 4 | 0 | 34,92 |
| Sbh1 | intra-protein › | 31 | 17 | 2131,121 | 711,381 | 3 | 0,85 | 34,89 |
| Sbh1 | intra-protein › | 17 | 15 | 1671,97 | 419 | 4 | 0,83 | 34,87 |
| Sbh1 | intra-protein › | 30 | 15 | 1681,024 | 421,264 | 4 | 0,67 | 34,31 |
| - | monolink | 41 | n/a | 3469,698 | 1157,574 | 3 | 0 | 34,23 |
| - | intralink | 23 | 30 | 1395,735 | 698,875 | 2 | 0 | 33,59 |
| - | intralink | 23 | 30 | 1523,83 | 762,923 | 2 | 0,59 | 33,26 |
| Sbh1 | intra-protein › | 31 | 17 | 2287,224 | 763,416 | 3 | 0,9 | 33,06 |
| - | monolink | 30 | n/a | 2012,09 | 1007,053 | 2 | 0,11 | 32,93 |
| Sbh1 | intra-protein › | 31 | 30 | 2140,187 | 536,055 | 4 | 0,71 | 32,67 |
| Sbh1 | intra-protein › | 31 | 23 | 3732,926 | 934,239 | 4 | 0,8 | 32,48 |
| - | monolink | 31 | n/a | 2012,091 | 671,705 | 3 | 0,11 | 31,81 |
| Sbh1 | intra-protein › | 41 | 23 | 3604,825 | 902,214 | 4 | 0 | 31,73 |
| Sbh1 | intra-protein › | 41 | 30 | 3245,726 | 650,153 | 5 | 0 | 31,56 |
| - | intralink | 30 | 41 | 1994,078 | 665,7 | 3 | 0,89 | 26,14 |

| *Crosslinked Peptide* |
|---|
| KQGSSQKVAASAPK-K7-156 |
| KNTNSNNSILKIYSDEATGLR-KQGSSQKVAASAPK-a11-b7 |
| MLISKSK-K5-156 |
| KQGSSQKVAASAPK-K7-155 |
| KQGSSQKVAASAPK-KQGSSQK-a7-b1 |
| KQGSSQKVAASAPK-K1-K7 |
| MLISKSK-K5-155 |
| VAASAPKK-K7-156 |
| NTNSNNSILKIYSDEATGLR-VAASAPKK-a10-b7 |
| VAASAPKK-K7-155 |
| KQGSSQKVAASAPK-KNTNSNNSILK-a7-b1 |
| VAASAPKK-K8-156 |
| MLISKSKMFK-K5-K7 |
| NTNSNNSILKIYSDEATGLR-K10-155 |
| VAASAPKK-K8-155 |
| KNTNSNNSILKIYSDEATGLR-K1-K11 |
| KNTNSNNSILKIYSDEATGLR-K11-156 |
| NTNSNNSILKIYSDEATGLR-K10-156 |
| QGSSQKVAASAPK-K6-156 |
| KNTNSNNSILK-K1-156 |
| KNTNSNNSILK-K1-155 |
| KNTNSNNSILKIYSDEATGLR-QGSSQKVAASAPK-a11-b6 |
| QGSSQKVAASAPK-KQGSSQK-a6-b1 |
| VAASAPKKNTNSNNSILK-K7-K8 |
| KNTNSNNSILKIYSDEATGLR-K11-155 |
| KQGSSQKVAASAPK-QGSSQKVAASAPK-a7-b6 |
| QGSSQKVAASAPK-TLQKRK-a6-b4 |
| QGSSQKVAASAPK-K6-155 |
| NTNSNNSILKIYSDEATGLRVDPENLYFQ-K10-156 |
| KQGSSQKVAASAPK-VAASAPKK-a7-b7 |
| KQGSSQK-K1-156 |
| MLISKSK-K5-K7 |
| NTNSNNSILKIYSDEATGLR-TLQKRK-a10-b4 |
| VAASAPKK-K7-K8 |
| KNTNSNNSILKIYSDEATGLR-RKQGSSQK-a11-b2 |
| KNTNSNNSILKIYSDEATGLR-KQGSSQK-a11-b1 |
| TLQKRK-K4-156 |
| NTNSNNSILKIYSDEATGLR-KQGSSQK-a10-b1 |
| VAASAPKK-KQGSSQK-a7-b1 |
| QGSSQKVAASAPK-KNTNSNNSILK-a6-b1 |
| KQGSSQKVAASAPK-K7-K14 |
| VAASAPKK-MLISKSK-a7-b5 |
| NTNSNNSILKIYSDEATGLR-KNTNSNNSILK-a10-b1 |
| QGSSQKVAASAPK-K6-K13 |
| RKQGSSQK-VAASAPKK-a2-b7 |
| KNTNSNNSILKIYSDEATGLR-K1-156 |
| KNTNSNNSILK-VAASAPKK-a1-b7 |
| KNTNSNNSILK-MLISKSK-a1-b5 |
| QGSSQKVAASAPKK-K6-156 |

NTNSNNNSILKIYSDEATGLR-QGSSQKVAASAPK-a10-b6
KQGSSQKVAASAPK-K1-156
KNTNSNNSILK-RKQGSSQK-a1-b2
KNTNSNNSILKIYSDEATGLR-NTNSNNSILKIYSDEATGLR-a11-b10
TLQKRK-K6-156
VAASAPKKNTNSNNSILK-K7-156
QGSSQKVAASAPKK-TLQKRK-a6-b4
KQGSSQK-TLQKRK-a1-b4
KNTNSNNSILKIYSDEATGLR-VAASAPKK-a11-b7
VAASAPKK-VAASAPKK-a7-b7
KNTNSNNSILK-KQGSSQK-a1-b1
VAASAPKKNTNSNNSILK-K8-156
MLISKSKMFK-K5-156
QGSSQKVAASAPKK-K6-K13
KQGSSQK-MLISKSK-a1-b5
NTNSNNSILKIYSDEATGLRVDPENLYFQ-K10-155
MLISKSK-K7-156
VAASAPKKNTNSNNSILK-K7-K18
MLISKSKMFK-K10-156
VAASAPKKNTNSNNSILK-K8-K18
MLISKSKMFK-K7-156

| ***Protein1*** | ***Protein2*** | ***XLType*** |
|---|---|---|
| Sbh1 | - | monolink |
| Sbh1 | Sbh1 | intra-protein › |
| Gls1 | - | monolink |
| Sbh1 | - | monolink |
| Sbh1 | Sbh1 | intra-protein › |
| Sbh1 | - | intralink |
| Gls1 | - | monolink |
| Sbh1 | - | monolink |
| Sbh1 | Sbh1 | intra-protein › |
| Sbh1 | - | monolink |
| Sbh1 | Sbh1 | intra-protein › |
| Sbh1 | - | monolink |
| Gls1 | - | intralink |
| Sbh1 | - | monolink |
| Sbh1 | - | monolink |
| Sbh1 | - | intralink |
| Sbh1 | - | monolink |
| Sbh1 | - | monolink |
| Sbh1 | - | monolink |
| Sbh1 | - | monolink |
| Sbh1 | - | monolink |
| Sbh1 | Sbh1 | intra-protein › |
| Sbh1 | Sbh1 | intra-protein › |
| Sbh1 | - | intralink |
| Sbh1 | - | monolink |
| Sbh1 | Sbh1 | intra-protein › |
| Sbh1 | Sbh1 | intra-protein › |
| Sbh1 | - | monolink |
| Sbh1 | - | monolink |
| Sbh1 | Sbh1 | intra-protein › |
| Sbh1 | - | monolink |
| Gls1 | - | intralink |
| Sbh1 | Sbh1 | intra-protein › |
| Sbh1 | - | intralink |
| Sbh1 | Sbh1 | intra-protein › |
| Sbh1 | Sbh1 | intra-protein › |
| Sbh1 | - | monolink |
| Sbh1 | Sbh1 | intra-protein › |
| Sbh1 | Sbh1 | intra-protein › |
| Sbh1 | Sbh1 | intra-protein › |
| Sbh1 | - | intralink |
| Sbh1 | Gls1 | inter-protein › |
| Sbh1 | Sbh1 | intra-protein › |
| Sbh1 | - | intralink |
| Sbh1 | Sbh1 | intra-protein › |
| Sbh1 | - | monolink |
| Sbh1 | Sbh1 | intra-protein › |
| Sbh1 | Gls1 | inter-protein › |
| Sbh1 | - | monolink |

| | | |
|---|---|---|
| Sbh1 | Sbh1 | intra-protein › |
| Sbh1 | - | monolink |
| Sbh1 | Sbh1 | intra-protein › |
| Sbh1 | Sbh1 | intra-protein › |
| Sbh1 | - | monolink |
| Sbh1 | - | monolink |
| Sbh1 | Sbh1 | intra-protein › |
| Sbh1 | Sbh1 | intra-protein › |
| Sbh1 | Sbh1 | intra-protein › |
| Sbh1 | Sbh1 | intra-protein › |
| Sbh1 | Sbh1 | intra-protein › |
| Sbh1 | - | monolink |
| Gls1 | - | monolink |
| Sbh1 | - | intralink |
| Sbh1 | Gls1 | inter-protein › |
| Sbh1 | - | monolink |
| Gls1 | - | monolink |
| Sbh1 | - | intralink |
| Gls1 | - | monolink |
| Sbh1 | - | intralink |
| Gls1 | - | monolink |

| Pos1 | Pos2 | Mr | Mz | z | deltaS | Id-Score |
|---|---|---|---|---|---|---|
| 23 | n/a | 1541,84 | 771,928 | 2 | 0,45 | 54,25 |
| 41 | 23 | 3861,024 | 644,512 | 6 | 0,75 | 52,86 |
| 5 | n/a | 961,549 | 481,783 | 2 | 0 | 52,65 |
| 23 | n/a | 1540,856 | 771,436 | 2 | 0,38 | 52,03 |
| 23 | 17 | 2285,232 | 572,316 | 4 | 0,74 | 49,96 |
| 17 | 23 | 1523,83 | 508,951 | 3 | 0,32 | 49,24 |
| 5 | n/a | 960,567 | 481,291 | 2 | 0 | 49,19 |
| 30 | n/a | 926,542 | 464,279 | 2 | 0,31 | 49,06 |
| 41 | 30 | 3117,632 | 780,416 | 4 | 0 | 48,7 |
| 30 | n/a | 925,559 | 463,788 | 2 | 0,33 | 47,89 |
| 23 | 31 | 2755,487 | 552,105 | 5 | 0,55 | 46,52 |
| 31 | n/a | 926,542 | 464,279 | 2 | 0 | 46,22 |
| 5 | 7 | 1349,745 | 675,88 | 2 | 0 | 45,74 |
| 41 | n/a | 2364,195 | 789,073 | 3 | 0 | 45,56 |
| 31 | n/a | 925,559 | 463,787 | 2 | 0,36 | 45,2 |
| 31 | 41 | 2475,254 | 826,092 | 3 | 0 | 45,19 |
| 41 | n/a | 2493,274 | 1247,645 | 2 | 0 | 45,1 |
| 41 | n/a | 2365,175 | 789,399 | 3 | 0 | 44,98 |
| 23 | n/a | 1413,745 | 707,881 | 2 | 0 | 44,63 |
| 31 | n/a | 1387,729 | 694,873 | 2 | 0 | 44,38 |
| 31 | n/a | 1386,744 | 694,38 | 2 | 0 | 44,14 |
| 41 | 23 | 3732,926 | 747,593 | 5 | 0,86 | 43,01 |
| 23 | 17 | 2157,141 | 720,055 | 3 | 0 | 42,57 |
| 30 | 31 | 1994,08 | 665,701 | 3 | 0 | 42,4 |
| 41 | n/a | 2492,283 | 1247,149 | 2 | 0 | 42,04 |
| 23 | 23 | 2781,499 | 696,383 | 4 | 0,85 | 41,63 |
| 23 | 15 | 2168,232 | 543,066 | 4 | 0 | 41,51 |
| 23 | n/a | 1412,762 | 707,389 | 2 | 0 | 41,39 |
| 41 | n/a | 3470,687 | 1157,904 | 3 | 0 | 41,12 |
| 23 | 30 | 2294,298 | 765,774 | 3 | 0,77 | 40,98 |
| 17 | n/a | 917,478 | 459,747 | 2 | 0 | 39,99 |
| 5 | 7 | 943,54 | 472,778 | 2 | 0 | 39,98 |
| 41 | 15 | 3119,658 | 624,939 | 5 | 0 | 39,2 |
| 30 | 31 | 908,532 | 455,274 | 2 | 0,59 | 39,09 |
| 41 | 17 | 3392,765 | 679,561 | 5 | 0,92 | 38,86 |
| 41 | 17 | 3236,662 | 810,173 | 4 | 0,35 | 38,63 |
| 15 | n/a | 928,57 | 465,293 | 2 | 0 | 38,6 |
| 41 | 17 | 3108,57 | 778,15 | 4 | 0 | 38,46 |
| 30 | 17 | 1669,935 | 557,653 | 3 | 0 | 38,1 |
| 23 | 31 | 2627,384 | 657,854 | 4 | 0 | 38,02 |
| 23 | 30 | 1523,83 | 762,923 | 2 | 0,48 | 38,01 |
| 30 | 5 | 1714,01 | 429,51 | 4 | 0,63 | 37,7 |
| 41 | 31 | 3578,814 | 895,711 | 4 | 0 | 37,57 |
| 23 | 30 | 1395,735 | 698,875 | 2 | 0 | 36,84 |
| 17 | 30 | 1826,036 | 609,687 | 3 | 0,75 | 36,8 |
| 31 | n/a | 2493,278 | 624,327 | 4 | 0 | 36,3 |
| 31 | 30 | 2140,186 | 714,403 | 3 | 0 | 36,23 |
| 31 | 5 | 2175,194 | 544,806 | 4 | 0 | 35,59 |
| 23 | n/a | 1541,841 | 514,955 | 3 | 0,55 | 35,5 |

| | | | | | | |
|---|---|---|---|---|---|---|
| 41 | 23 | 3604,842 | 902,218 | 4 | 0 | 35,04 |
| 17 | n/a | 1541,842 | 514,955 | 3 | 0,33 | 34,59 |
| 31 | 17 | 2287,222 | 763,415 | 3 | 0 | 34,57 |
| 41 | 41 | 4684,355 | 937,879 | 5 | 0,29 | 34,35 |
| 17 | n/a | 928,568 | 465,292 | 2 | 0 | 34,24 |
| 30 | n/a | 2012,09 | 671,704 | 3 | 0 | 34,11 |
| 23 | 15 | 2296,321 | 460,272 | 5 | 0,7 | 33,99 |
| 17 | 15 | 1671,97 | 419 | 4 | 0 | 33,66 |
| 41 | 30 | 3245,728 | 650,153 | 5 | 0 | 33,45 |
| 30 | 30 | 1678,994 | 420,756 | 4 | 0,78 | 33,3 |
| 31 | 17 | 2131,12 | 711,381 | 3 | 0 | 33,16 |
| 31 | n/a | 2012,091 | 671,705 | 3 | 0,15 | 33,14 |
| 5 | n/a | 1367,755 | 684,886 | 2 | 0 | 33,14 |
| 23 | 30 | 1523,829 | 762,922 | 2 | 0,59 | 32,63 |
| 17 | 5 | 1704,948 | 427,245 | 4 | 0 | 30,25 |
| 41 | n/a | 3469,701 | 1157,575 | 3 | 0 | 28,78 |
| 7 | n/a | 961,551 | 481,783 | 2 | 0 | 28,02 |
| 30 | 41 | 1994,083 | 665,702 | 3 | 0,24 | 27,83 |
| 10 | n/a | 1367,755 | 684,885 | 2 | 0 | 26,84 |
| 31 | 41 | 1994,079 | 665,701 | 3 | 0,41 | 26,18 |
| 7 | n/a | 1367,754 | 456,926 | 3 | 0 | 25,37 |

| *Crosslinked Peptide* |
|---|
| KQGSSQKVAASAPK-K7-156 |
| KQGSSQKVAASAPK-K7-155 |
| KNTNSNNSILKIYSDEATGLR-KQGSSQKVAASAPK-a11-b7 |
| VAASAPKK-K7-156 |
| KQGSSQKVAASAPK-K1-K7 |
| KQGSSQKVAASAPK-KNTNSNNSILK-a7-b1 |
| VAASAPKK-K8-156 |
| KQGSSQKVAASAPK-QGSSQKVAASAPK-a7-b6 |
| KQGSSQKVAASAPK-KQGSSQK-a7-b1 |
| VAASAPKK-K8-155 |
| NTNSNNSILKIYSDEATGLR-K10-156 |
| NTNSNNSILKIYSDEATGLR-VAASAPKK-a10-b7 |
| KNTNSNNSILKIYSDEATGLR-K1-K11 |
| KNTNSNNSILKIYSDEATGLR-K11-156 |
| NTNSNNSILKIYSDEATGLR-K10-155 |
| VAASAPKK-K7-155 |
| KNTNSNNSILKIYSDEATGLR-KNTNSNNSILK-a11-b1 |
| QGSSQKVAASAPK-K6-156 |
| KNTNSNNSILK-K1-156 |
| KNTNSNNSILKIYSDEATGLR-KQGSSQK-a11-b1 |
| KNTNSNNSILKIYSDEATGLR-K11-155 |
| KNTNSNNSILK-K1-155 |
| KNTNSNNSILKIYSDEATGLR-QGSSQKVAASAPK-a11-b6 |
| KNTNSNNSILKIYSDEATGLR-VAASAPKK-a11-b7 |
| KNTNSNNSILK-KNTNSNNSILK-a1-b1 |
| KQGSSQK-K1-156 |
| QGSSQKVAASAPK-K6-155 |
| KNTNSNNSILK-K11-156 |
| NTNSNNSILKIYSDEATGLR-TLQKRK-a10-b4 |
| VAASAPKKNTNSNNSILK-K7-K8 |
| TLQKRK-K4-156 |
| QGSSQKVAASAPK-TLQKRK-a6-b4 |
| KQGSSQKVAASAPK-VAASAPKK-a7-b7 |
| QGSSQKVAASAPKK-TLQKRK-a6-b4 |
| NTNSNNSILKIYSDEATGLRVDPENLYFQ-K10-156 |
| QGSSQKVAASAPK-KQGSSQK-a6-b1 |
| VAASAPKK-K7-K8 |
| VAASAPKKNTNSNNSILK-K7-156 |
| VAASAPKK-KQGSSQK-a7-b1 |
| QGSSQKVAASAPKK-K6-156 |
| NTNSNNSILKIYSDEATGLR-KNTNSNNSILK-a10-b1 |
| RKQGSSQK-VAASAPKK-a2-b7 |
| NTNSNNSILKIYSDEATGLR-KQGSSQK-a10-b1 |
| VAASAPKK-VAASAPKK-a7-b7 |
| QGSSQKVAASAPK-KNTNSNNSILK-a6-b1 |
| QGSSQKVAASAPK-VAASAPKK-a6-b7 |
| VAASAPKK-TLQKRK-a7-b4 |
| KQGSSQKVAASAPK-K7-K14 |
| QGSSQKVAASAPKK-K6-K13 |

KNTNSNNSILKIYSDEATGLR-K1-156
NTNSNNSILKIYSDEATGLRVDPENLYFQ-K10-155
VAASAPKKNTNSNNSILK-K8-156
QGSSQKVAASAPK-K6-K13

| *Protein1* | *Protein2* | *XLType* |
|---|---|---|
| Sbh1 | - | monolink |
| Sbh1 | - | monolink |
| Sbh1 | Sbh1 | intra-protein › |
| Sbh1 | - | monolink |
| Sbh1 | - | intralink |
| Sbh1 | Sbh1 | intra-protein › |
| Sbh1 | - | monolink |
| Sbh1 | Sbh1 | intra-protein › |
| Sbh1 | Sbh1 | intra-protein › |
| Sbh1 | - | monolink |
| Sbh1 | - | monolink |
| Sbh1 | Sbh1 | intra-protein › |
| Sbh1 | - | intralink |
| Sbh1 | - | monolink |
| Sbh1 | - | monolink |
| Sbh1 | - | monolink |
| Sbh1 | Sbh1 | intra-protein › |
| Sbh1 | - | monolink |
| Sbh1 | - | monolink |
| Sbh1 | Sbh1 | intra-protein › |
| Sbh1 | - | monolink |
| Sbh1 | - | monolink |
| Sbh1 | Sbh1 | intra-protein › |
| Sbh1 | Sbh1 | intra-protein › |
| Sbh1 | Sbh1 | intra-protein › |
| Sbh1 | - | monolink |
| Sbh1 | - | monolink |
| Sbh1 | - | monolink |
| Sbh1 | Sbh1 | intra-protein › |
| Sbh1 | - | intralink |
| Sbh1 | - | monolink |
| Sbh1 | Sbh1 | intra-protein › |
| Sbh1 | Sbh1 | intra-protein › |
| Sbh1 | Sbh1 | intra-protein › |
| Sbh1 | - | monolink |
| Sbh1 | Sbh1 | intra-protein › |
| Sbh1 | - | intralink |
| Sbh1 | - | monolink |
| Sbh1 | Sbh1 | intra-protein › |
| Sbh1 | - | monolink |
| Sbh1 | Sbh1 | intra-protein › |
| Sbh1 | Sbh1 | intra-protein › |
| Sbh1 | Sbh1 | intra-protein › |
| Sbh1 | Sbh1 | intra-protein › |
| Sbh1 | Sbh1 | intra-protein › |
| Sbh1 | Sbh1 | intra-protein › |
| Sbh1 | Sbh1 | intra-protein › |
| Sbh1 | - | intralink |
| Sbh1 | - | intralink |

| | | |
|---|---|---|
| Sbh1 | - | monolink |
| Sbh1 | - | monolink |
| Sbh1 | - | monolink |
| Sbh1 | - | intralink |

| Pos1 | Pos2 | Mr | Mz | z | deltaS | Id-Score |
|---|---|---|---|---|---|---|
| 23 | n/a | 1541,84 | 771,928 | 2 | 0,42 | 53,61 |
| 23 | n/a | 1540,856 | 771,436 | 2 | 0,34 | 52,07 |
| 41 | 23 | 3861,028 | 644,512 | 6 | 0,68 | 49,67 |
| 30 | n/a | 926,542 | 464,279 | 2 | 0,58 | 47,2 |
| 17 | 23 | 1523,828 | 508,95 | 3 | 0,14 | 47,1 |
| 23 | 31 | 2755,483 | 552,104 | 5 | 0,55 | 46,93 |
| 31 | n/a | 926,542 | 464,279 | 2 | 0,58 | 46,86 |
| 23 | 23 | 2781,5 | 557,308 | 5 | 0,89 | 46,82 |
| 23 | 17 | 2285,235 | 572,316 | 4 | 0,71 | 46,76 |
| 31 | n/a | 925,559 | 463,787 | 2 | 0,56 | 46,25 |
| 41 | n/a | 2365,176 | 789,4 | 3 | 0 | 45,54 |
| 41 | 30 | 3117,628 | 780,415 | 4 | 0,77 | 45,3 |
| 31 | 41 | 2475,26 | 826,095 | 3 | 0 | 45,25 |
| 41 | n/a | 2493,269 | 1247,642 | 2 | 0 | 44,84 |
| 41 | n/a | 2364,194 | 789,072 | 3 | 0 | 44,84 |
| 30 | n/a | 925,559 | 463,787 | 2 | 0,57 | 44,25 |
| 41 | 31 | 3706,91 | 742,39 | 5 | 0 | 43,64 |
| 23 | n/a | 1413,743 | 472,255 | 3 | 0 | 43,22 |
| 31 | n/a | 1387,729 | 694,872 | 2 | 0 | 43,1 |
| 41 | 17 | 3236,666 | 810,174 | 4 | 0,34 | 42,89 |
| 41 | n/a | 2492,288 | 624,08 | 4 | 0 | 42,15 |
| 31 | n/a | 1386,745 | 694,38 | 2 | 0 | 41,91 |
| 41 | 23 | 3732,93 | 934,24 | 4 | 0,78 | 41,79 |
| 41 | 30 | 3245,726 | 812,439 | 4 | 0,78 | 41,65 |
| 31 | 31 | 2601,37 | 868,131 | 3 | 0 | 41,6 |
| 17 | n/a | 917,479 | 459,747 | 2 | 0 | 41,14 |
| 23 | n/a | 1412,762 | 707,389 | 2 | 0 | 41,07 |
| 41 | n/a | 1387,731 | 694,873 | 2 | 0 | 40,82 |
| 41 | 15 | 3119,656 | 624,939 | 5 | 0,13 | 40,76 |
| 30 | 31 | 1994,08 | 665,701 | 3 | 0,2 | 40,58 |
| 15 | n/a | 928,57 | 465,293 | 2 | 0 | 40,16 |
| 23 | 15 | 2168,223 | 434,652 | 5 | 0 | 40,02 |
| 23 | 30 | 2294,297 | 765,773 | 3 | 0,9 | 39,88 |
| 23 | 15 | 2296,32 | 460,272 | 5 | 0,85 | 39,74 |
| 41 | n/a | 3470,691 | 1157,905 | 3 | 0 | 39,73 |
| 23 | 17 | 2157,14 | 720,054 | 3 | 0 | 39,7 |
| 30 | 31 | 908,532 | 455,274 | 2 | 0,49 | 39,22 |
| 30 | n/a | 2012,091 | 671,705 | 3 | 0 | 37,8 |
| 30 | 17 | 1669,935 | 557,653 | 3 | 0,82 | 37,52 |
| 23 | n/a | 1541,842 | 514,955 | 3 | 0,39 | 36,93 |
| 41 | 31 | 3578,819 | 1193,948 | 3 | 0 | 36,78 |
| 17 | 30 | 1826,035 | 609,686 | 3 | 0,87 | 36,65 |
| 41 | 17 | 3108,572 | 778,151 | 4 | 0 | 36,34 |
| 30 | 30 | 1679,001 | 420,758 | 4 | 0,85 | 36,1 |
| 23 | 31 | 2627,385 | 657,854 | 4 | 0,26 | 36,01 |
| 23 | 30 | 2166,2 | 723,074 | 3 | 0,95 | 35,63 |
| 30 | 15 | 1681,029 | 421,265 | 4 | 0,72 | 35,21 |
| 23 | 30 | 1523,831 | 762,923 | 2 | 0,66 | 33,83 |
| 23 | 30 | 1523,829 | 762,922 | 2 | 0,56 | 33,81 |

| | | | | | | |
|---|---|---|---|---|---|---|
| 31 | n/a | 2493,274 | 1247,645 | 2 | 0 | 33,03 |
| 41 | n/a | 3469,708 | 1157,577 | 3 | 0 | 32,26 |
| 31 | n/a | 2012,091 | 1007,054 | 2 | 0,11 | 28,91 |
| 23 | 30 | 1395,735 | 698,875 | 2 | 0 | 28,14 |

| *Crosslinked Peptide* |
|---|
| KQGSSQKVAASAPK-K7-156 |
| KQGSSQKVAASAPK-K1-K7 |
| NTNSNNSILKIYSDEATGLR-VAASAPKK-a10-b7 |
| VAASAPKK-K8-156 |
| KQGSSQKVAASAPK-K7-155 |
| KQGSSQKVAASAPK-KNTNSNNSILK-a7-b1 |
| VAASAPKK-K7-156 |
| KNTNSNNSILKIYSDEATGLR-K11-156 |
| KNTNSNNSILKIYSDEATGLR-K1-K11 |
| VAASAPKK-K7-155 |
| VAASAPKK-K8-155 |
| KNTNSNNSILK-K1-155 |
| QGSSQKVAASAPK-K6-156 |
| QGSSQKVAASAPK-TLQKRK-a6-b4 |
| KNTNSNNSILK-K1-156 |
| KQGSSQKVAASAPK-VAASAPKK-a7-b7 |
| KNTNSNNSILKIYSDEATGLR-KQGSSQK-a11-b1 |
| NTNSNNSILKIYSDEATGLR-K10-155 |
| KNTNSNNSILKIYSDEATGLR-KNTNSNNSILK-a11-b1 |
| NTNSNNSILKIYSDEATGLR-K10-156 |
| KQGSSQKVAASAPK-KQGSSQK-a7-b1 |
| KNTNSNNSILKIYSDEATGLR-K11-155 |
| KNTNSNNSILKIYSDEATGLR-QGSSQKVAASAPK-a11-b6 |
| VAASAPKK-K7-K8 |
| VAASAPKKNTNSNNSILK-K7-K8 |
| VAASAPKK-KQGSSQK-a7-b1 |
| QGSSQKVAASAPK-KQGSSQK-a6-b1 |
| TLQKRK-K4-156 |
| NTNSNNSILKIYSDEATGLRVDPENLYFQ-K10-156 |
| KNTNSNNSILKIYSDEATGLR-VAASAPKK-a11-b7 |
| KQGSSQK-K1-156 |
| QGSSQKVAASAPK-K6-155 |
| VAASAPKK-VAASAPKK-a7-b7 |
| NTNSNNSILKIYSDEATGLR-KQGSSQK-a10-b1 |
| QGSSQKVAASAPKK-K6-156 |
| NTNSNNSILKIYSDEATGLR-TLQKRK-a10-b4 |
| KQGSSQK-TLQKRK-a1-b4 |
| VAASAPKKNTNSNNSILK-KQGSSQK-a7-b1 |
| QGSSQKVAASAPKK-TLQKRK-a6-b4 |
| KNTNSNNSILK-KNTNSNNSILK-a1-b1 |
| KQGSSQKVAASAPK-QGSSQKVAASAPK-a7-b6 |
| KNTNSNNSILKIYSDEATGLR-NTNSNNSILKIYSDEATGLR-a11-b10 |
| RKQGSSQK-VAASAPKK-a2-b7 |
| QGSSQKVAASAPK-KNTNSNNSILK-a6-b1 |
| KNTNSNNSILK-VAASAPKK-a1-b7 |
| VAASAPKKNTNSNNSILK-K7-156 |
| KNTNSNNSILK-KQGSSQK-a1-b1 |
| KNTNSNNSILKIYSDEATGLR-KQGSSQKVAASAPK-a11-b7 |
| VAASAPKK-TLQKRK-a7-b4 |

QGSSQKVAASAPKK-K6-K13
NTNSNNSILKIYSDEATGLR-RKQGSSQK-a10-b2
KNTNSNNSILKIYSDEATGLR-K1-156
QGSSQKVAASAPK-K6-K13
NTNSNNSILKIYSDEATGLRVDPENLYFQ-K10-155
KQGSSQK-K1-K7
KQGSSQKVAASAPK-K7-K14
VAASAPKKNTNSNNSILK-K8-K18
KQGSSQKVAASAPK-K1-156
QGSSQKVAASAPKK-K6-K14
MKNSVGISIATIVAIIAAX-QGSSQKVAASAPK-a2-b6

| ***Protein1*** | ***Protein2*** | ***XLType*** |
|---|---|---|
| Sbh1 | - | monolink |
| Sbh1 | - | intralink |
| Sbh1 | Sbh1 | intra-protein › |
| Sbh1 | - | monolink |
| Sbh1 | - | monolink |
| Sbh1 | Sbh1 | intra-protein › |
| Sbh1 | - | monolink |
| Sbh1 | - | monolink |
| Sbh1 | - | intralink |
| Sbh1 | - | monolink |
| Sbh1 | - | monolink |
| Sbh1 | - | monolink |
| Sbh1 | - | monolink |
| Sbh1 | Sbh1 | intra-protein › |
| Sbh1 | - | monolink |
| Sbh1 | Sbh1 | intra-protein › |
| Sbh1 | Sbh1 | intra-protein › |
| Sbh1 | - | monolink |
| Sbh1 | Sbh1 | intra-protein › |
| Sbh1 | - | monolink |
| Sbh1 | Sbh1 | intra-protein › |
| Sbh1 | - | monolink |
| Sbh1 | Sbh1 | intra-protein › |
| Sbh1 | - | intralink |
| Sbh1 | - | intralink |
| Sbh1 | Sbh1 | intra-protein › |
| Sbh1 | Sbh1 | intra-protein › |
| Sbh1 | - | monolink |
| Sbh1 | - | monolink |
| Sbh1 | Sbh1 | intra-protein › |
| Sbh1 | - | monolink |
| Sbh1 | - | monolink |
| Sbh1 | Sbh1 | intra-protein › |
| Sbh1 | Sbh1 | intra-protein › |
| Sbh1 | - | monolink |
| Sbh1 | Sbh1 | intra-protein › |
| Sbh1 | Sbh1 | intra-protein › |
| Sbh1 | Sbh1 | intra-protein › |
| Sbh1 | Sbh1 | intra-protein › |
| Sbh1 | Sbh1 | intra-protein › |
| Sbh1 | Sbh1 | intra-protein › |
| Sbh1 | Sbh1 | intra-protein › |
| Sbh1 | Sbh1 | intra-protein › |
| Sbh1 | Sbh1 | intra-protein › |
| Sbh1 | Sbh1 | intra-protein › |
| Sbh1 | - | monolink |
| Sbh1 | Sbh1 | intra-protein › |
| Sbh1 | Sbh1 | intra-protein › |
| Sbh1 | Sbh1 | intra-protein › |

| | | |
|---|---|---|
| Sbh1 | - | intralink |
| Sbh1 | Sbh1 | intra-protein x |
| Sbh1 | - | monolink |
| Sbh1 | - | intralink |
| Sbh1 | - | monolink |
| Sbh1 | - | intralink |
| Sbh1 | - | intralink |
| Sbh1 | - | intralink |
| Sbh1 | - | monolink |
| Sbh1 | - | intralink |
| Mns1 | Sbh1 | inter-protein x |

| *Pos1* | *Pos2* | *Mr* | *Mz* | *z* | *deltaS* | *Id-Score* |
|---|---|---|---|---|---|---|
| 23 | n/a | 1541,839 | 771,927 | 2 | 0,4 | 52,51 |
| 17 | 23 | 1523,83 | 508,951 | 3 | 0,32 | 48,43 |
| 41 | 30 | 3117,628 | 780,415 | 4 | 0,77 | 48,21 |
| 31 | n/a | 926,543 | 464,279 | 2 | 0,43 | 47,75 |
| 23 | n/a | 1540,859 | 771,437 | 2 | 0,41 | 47,74 |
| 23 | 31 | 2755,483 | 552,104 | 5 | 0,67 | 46,88 |
| 30 | n/a | 926,543 | 464,279 | 2 | 0,46 | 46,67 |
| 41 | n/a | 2493,268 | 832,097 | 3 | 0 | 45,84 |
| 31 | 41 | 2475,261 | 826,095 | 3 | 0 | 45,68 |
| 30 | n/a | 925,557 | 463,786 | 2 | 0,44 | 45,1 |
| 31 | n/a | 925,558 | 463,787 | 2 | 0,47 | 43,79 |
| 31 | n/a | 1386,743 | 694,379 | 2 | 0 | 43,69 |
| 23 | n/a | 1413,744 | 707,88 | 2 | 0 | 43,14 |
| 23 | 15 | 2168,229 | 434,654 | 5 | 0 | 42,85 |
| 31 | n/a | 1387,729 | 694,872 | 2 | 0 | 42,78 |
| 23 | 30 | 2294,293 | 459,866 | 5 | 0,87 | 42,59 |
| 41 | 17 | 3236,661 | 810,173 | 4 | 0,31 | 42,05 |
| 41 | n/a | 2364,196 | 1183,106 | 2 | 0 | 41,92 |
| 41 | 31 | 3706,916 | 927,737 | 4 | 0 | 41,62 |
| 41 | n/a | 2365,179 | 789,401 | 3 | 0 | 41,52 |
| 23 | 17 | 2285,23 | 762,751 | 3 | 0,7 | 41,33 |
| 41 | n/a | 2492,281 | 1247,148 | 2 | 0 | 41,02 |
| 41 | 23 | 3732,931 | 934,241 | 4 | 0,73 | 40,93 |
| 30 | 31 | 908,531 | 455,274 | 2 | 0,43 | 40,48 |
| 30 | 31 | 1994,081 | 665,701 | 3 | 0,06 | 40,08 |
| 30 | 17 | 1669,934 | 557,652 | 3 | 0,61 | 39,65 |
| 23 | 17 | 2157,141 | 720,055 | 3 | 0 | 39,07 |
| 15 | n/a | 928,57 | 465,293 | 2 | 0 | 38,83 |
| 41 | n/a | 3470,691 | 1157,905 | 3 | 0 | 38,63 |
| 41 | 30 | 3245,726 | 650,153 | 5 | 0 | 38,24 |
| 17 | n/a | 917,481 | 459,748 | 2 | 0 | 37,84 |
| 23 | n/a | 1412,761 | 707,388 | 2 | 0 | 37,74 |
| 30 | 30 | 1678,996 | 420,757 | 4 | 0,92 | 37,56 |
| 41 | 17 | 3108,572 | 1037,198 | 3 | 0 | 37,51 |
| 23 | n/a | 1541,842 | 514,955 | 3 | 0,54 | 37,22 |
| 41 | 15 | 3119,655 | 624,939 | 5 | 0,21 | 37 |
| 17 | 15 | 1671,968 | 419 | 4 | 0 | 36,73 |
| 30 | 17 | 2755,484 | 689,879 | 4 | 0,83 | 36,08 |
| 23 | 15 | 2296,32 | 460,272 | 5 | 0,8 | 36,06 |
| 31 | 31 | 2601,377 | 1301,696 | 2 | 0 | 35,87 |
| 23 | 23 | 2781,496 | 696,382 | 4 | 0,9 | 35,04 |
| 41 | 41 | 4684,355 | 1172,097 | 4 | 0 | 34,99 |
| 17 | 30 | 1826,036 | 609,686 | 3 | 0,8 | 34,59 |
| 23 | 31 | 2627,389 | 876,804 | 3 | 0 | 34,48 |
| 31 | 30 | 2140,183 | 714,402 | 3 | 0,66 | 34,38 |
| 30 | n/a | 2012,09 | 1007,053 | 2 | 0,19 | 34,28 |
| 31 | 17 | 2131,121 | 711,381 | 3 | 0 | 34,2 |
| 41 | 23 | 3861,023 | 773,212 | 5 | 0,74 | 33,89 |
| 30 | 15 | 1681,028 | 421,265 | 4 | 0,66 | 33,73 |

| | | | | | | |
|---|---|---|---|---|---|---|
| 23 | 30 | 1523,831 | 762,923 | 2 | 0,52 | 33,62 |
| 41 | 17 | 3264,661 | 817,173 | 4 | 0 | 33,32 |
| 31 | n/a | 2493,273 | 1247,645 | 2 | 0 | 33,08 |
| 23 | 30 | 1395,735 | 698,875 | 2 | 0 | 29,18 |
| 41 | n/a | 3469,694 | 1157,573 | 3 | 0 | 26,7 |
| 17 | 23 | 899,469 | 450,742 | 2 | 0 | 24,96 |
| 23 | 30 | 1523,83 | 762,923 | 2 | 0,45 | 24,29 |
| 31 | 41 | 1994,082 | 998,049 | 2 | 0,23 | 22,65 |
| 17 | n/a | 1541,841 | 514,955 | 3 | 0,77 | 21,99 |
| 23 | 31 | 1523,83 | 762,923 | 2 | 0,73 | 21,34 |
| 2 | 23 | 3520,926 | 705,193 | 5 | 0,87 | 20,6 |
